# Evolutionary Origins of Molecular Programs Underlying Brain Circuitry

**DOI:** 10.64898/2026.06.29.735061

**Authors:** Zhi Huang, Shenglong Li, Zhenkun Zhuang, Duoyuan Chen, Xiao Du, Qun Liu, Cirong Liu, Shaojie Ma, Yidi Sun, Hongli Du, Shiping Liu, Guangyi Fan, Longqi Liu, Shijie Hao

## Abstract

The vertebrate pallium harbors independently evolved structures that, nevertheless, support strikingly similar sensory and cognitive circuit architectures. The mechanisms and evolutionary timing driving the emergence of these parallel pallial circuits remain unelucidated. Here, we integrated spatial transcriptomic and single-nucleus RNA-seq datasets from eight representative vertebrate species spanning approximately 500 million years of evolution to reconstruct the evolutionary assembly of the primary sensory-allocortical (Pr-Al) molecular axis, a conserved cortical hierarchical axis defined in our prior work. We uncovered that Pr- and Al-like neuronal identities are deeply conserved across sarcopterygians, encompassing all tetrapods and lobe-finned fishes. Intriguingly, this ancestral neuronal homology is uncoupled from spatially partitioned patterning: only tetrapods further compartmentalized these Pr- and Al- neurons into distinct pallium regions. Functional enrichment of Pr- and Al- gene programs uncovered a conserved tetrapod genetic core suite including MAPK signaling and axon guidance pathways. This core toolkit underwent sequential functional refinement from amphibians through mammals. Notably, mammals and birds convergently evolved association-cortical molecular profiles enriched for synaptic regulatory genes to support advanced cognitive functions. Collectively, this work delineates a stepwise vertebrate pallium evolutionary paradigm that explains how conserved molecular modules shape spatially organized brain circuitry across deep evolutionary time.

## Introduction

The vertebrate pallium has undergone profound anatomical diversification, yet exhibits strikingly conserved principles of sensory processing. This contrast is most evident in the relationship between the mammalian neocortex and the sauropsid dorsal ventricular ridge (DVR). The neocortex is laminated and derives from the dorsal pallium, whereas the DVR is organized as nuclear structures and derives primarily from the ventral pallium. How these distinct pallial architectures came to support comparable input-output sensory circuits remains one of the central questions in comparative neurobiology^1–3^.

Historically, this question was framed as a debate over structural homology. The recapitulation hypothesis proposed that a DVR-like ancestral structure was retained in sauropsids and transformed into the mammalian neocortex, whereas the out-group hypothesis argued that the neocortex and DVR evolved independently as parallel pallial adaptations^4–6^. Developmental fate mapping and the tetrapartite pallium model later established a broad structural consensus: the amniote pallium contains conserved medial, dorsal, lateral, and ventral domains, and the neocortex and DVR originate from different pallial territories^7, 8^. Thus, the strict structural version of DVR-neocortex homology has largely been rejected.

This structural resolution, however, did not resolve the deeper biological paradox. Despite their distinct developmental origins, the DVR and neocortex share multiple features of circuit organization, including related neuronal molecular profiles, comparable input and output cell classes, topologically organized thalamic projections, and hierarchical sensory processing architectures^9–19^. These observations suggest that vertebrate pallia may deploy conserved cellular and molecular programs to build similar circuit logic in non-homologous anatomical substrates. Although previous studies on sauropsids have indicated links between genetic programs and neural circuits, such evidence has so far been limited to the avian vocal learning circuit^20^. A framework based solely on regional homology is therefore insufficient; what is needed is a quantitative axis that can compare circuit organization across divergent pallial structures.

The primary sensory-allocortical (Pr-Al) axis provides such a framework. Originally defined in mammalian cortex, the Pr-Al index describes opposing molecular programs anchored by primary sensory regions at the Pr pole and allocortical or periallocortical regions at the Al pole^21–23^. This axis tracks the flow of pallial information processing from sensory input regions, through association territories, toward output-related allocortical regions. It is also associated with distinct neuronal populations, gene modules, thalamocortical connectivity patterns, and functional hierarchy (**Fig. 1A**). The Pr-Al index therefore offers a way to ask whether structurally different pallia share a common molecular logic for organizing sensory-to-output circuits.

**Figure 1.**
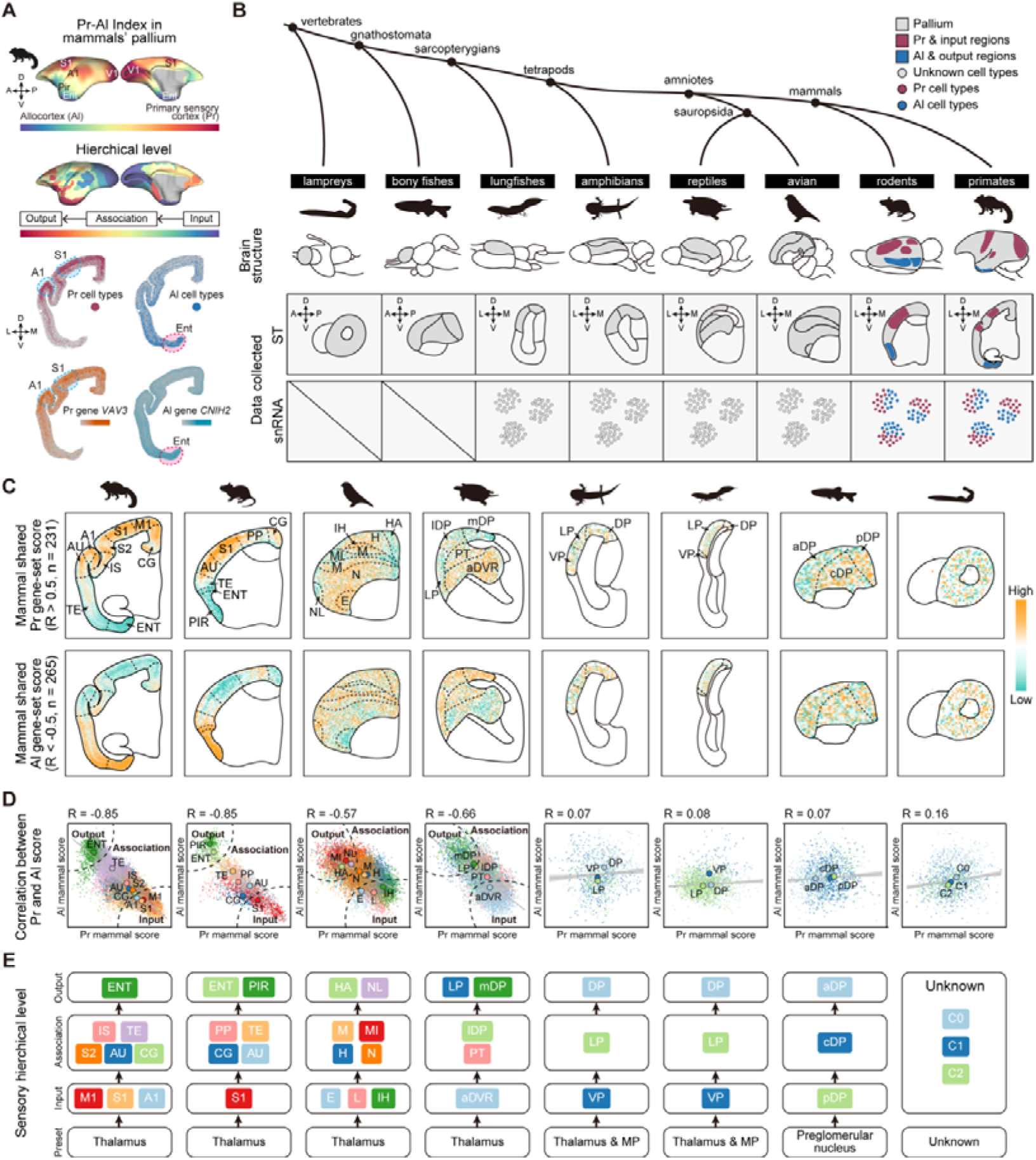
Evolutionary conservation analysis of the Pr-Al index across vertebrates. **(A)** Top panel depicts the Pr-Al index representation in the marmoset cortex, showing a continuous molecular gradient from primary sensory cortex (Pr, red) to allocortex (Al, blue)^21^. Middle panel illustrates the corresponding cortical hierarchical level gradient in marmoset^36^. Bottom spatial plots demonstrate the expression patterns of Pr- and Al-associated genes and their respective cell populations. **(B)** Phylogenetic tree of vertebrate evolution, including eight species representing all major vertebrate clades. Schematic brain sections illustrate pallial anatomy, and collected data types are indicated (ST: spatial transcriptomics; snRNA-seq: single-nucleus RNA sequencing). Notably, the schematics for mouse and marmoset additionally delineate the anatomical positions of Pr and Al regions. **(C)** Spatial plots of the mammalian-derived conserved Pr gene set (n=231, Pearson R > 0.5 with Pr index) and Al gene set (n=265, Pearson R < −0.5 with Al index) across all species. **(D)** Scatter plots showing Pearson correlation coefficients (R) between Pr and Al gene set scores for each species. Dash lines indicate the boundaries of output, association and input regions. **(E)** Reconstructed pallium sensory processing hierarchies across vertebrates, see Methods for details. Region abbreviations see **fig. S1**.

Several key evolutionary questions remain unresolved. First, it is unknown whether a Pr-Al index exists outside mammals, particularly in sauropsid pallial regions such as the DVR that are functionally analogous but not structurally homologous to neocortex. Second, if Pr-Al index is conserved, it is unclear whether conservation occurs first at the level of neuronal identity, spatial tissue organization, or both. Third, the timing and molecular mechanisms by which this axis became linked to sensory hierarchies and advanced association pallial functions remain poorly understood.

Here, we address these questions by integrating spatial transcriptomic and single-nucleus RNA-seq datasets across eight representative vertebrate species, including the basal jawless vertebrate sea lamprey (*Petromyzon marinus*), the ray-finned fish zebrafish (*Danio rerio*), the lobe-finned fish West African lungfish (*Protopterus annectens*), the amphibian Mexican axolotl (*Ambystoma mexicanum*), the reptile Chinese softshell turtle (*Pelodiscus sinensis*), the avian zebra finch (*Taeniopygia guttata*), the rodent house mouse (*Mus musculus*), and the primate common marmoset (*Callithrix jacchus*). Using cross-species orthogroup mapping, Pr-Al gene-set scoring, neuronal subtype integration, spatial deconvolution, and comparative genomic analyses, we reconstruct the evolutionary assembly of the Pr-Al molecular axis across approximately 500 million years of vertebrate evolution. Our results reveal a stepwise process in which ancient Pr- and Al-like neuronal identities predated tetrapods, were spatially organized into a Pr-Al axis during tetrapod evolution, and were later refined in amniotes and independently elaborated in mammals and birds. This model reconciles the absence of strict structural homology between the neocortex and DVR with their shared circuit-level organization, and provides a molecular framework for understanding the parallel evolution of complex pallial functions.

## Results

### Mammalian-conserved Pr-Al gene programs reveal a conserved pallial input-output axis in amniotes

To systematically explore the distribution and evolutionary origin of the Pr-Al index across vertebrates, we integrated spatially resolved transcriptomic (ST) and single-nucleus RNA sequencing (snRNA-seq) datasets from eight key vertebrate evolutionary nodes, covering approximately 500 million years of evolution (**Fig. 1B**). Our sampling covered all major vertebrate clades: jawless vertebrates (lamprey)^24^, ray-finned fishes (zebrafish), lobe-finned fishes (lungfish), Amphibia (axolotl)^25,26^, sauropsids (Chinese softshell turtle^27^, zebra finch^27^), Mammalia (rodent: mouse^28^, primate: common marmoset^21^).

First, we manually delineated the pallial regions across all species based on their respective anatomical annotations, and explicitly separated the hippocampus and its homologous structure, the medial pallium (MP)^9^. The MP was excluded from subsequent Pr-Al index analyses owing to its distinct connectivity patterns, structural characteristics and cellular composition^29^. We next performed unsupervised clustering independently on ST data from each species, and identified brain regions that were highly concordant with anatomical partitions, including those of avian^30^ (**fig. S1A-B**), turtle^31^ (**fig. S1C-D**), axolotl^32^ (**fig. S1E-F**), lungfish^33^ (**fig. S1G-H**). and zebrafish^10^ (**fig. S1I-J**). Of note, no spatially distinct brain regions were reported previously or detected in our analysis in the lamprey pallium, thus we used clusters instead of brain regions for further analysis (**fig. S1K**). Pallial partitions for the mouse and marmoset were defined based on standard published brain atlas annotations^28^ (**fig. S1L**).

We next established a unified cross-species orthogroup framework using OrthoFinder and projected gene expression from each species onto shared orthogroups (**fig. S2A–B**)^34^. Based on previously defined Pr-Al profiles in mouse and marmoset, we identified mammalian-conserved orthogroups that were positively associated with the Pr pole or negatively associated with the Al pole. These Pr- and Al-related gene sets were then used to score spatial transcriptomic datasets across all examined species.

Using this mammalian-conserved Pr-Al gene program, we detected robust antagonistic Pr and Al spatial patterns in all amniote species examined, including mammals, avian, and turtle. In these species, Pr-related genes were preferentially enriched in sensory input regions, whereas Al-related genes were enriched in output-related pallial regions, with association territories occupying intermediate positions **(Fig. 1C-E, see Methods)**^15, 18, 30^. Specifically, in marmosets, Pr/input signals were highest in primary sensory areas S1, M1, and A1, whereas Al/output signals in the entorhinal cortex (ENT), a major output region. In mice, Pr/input signals were detected in S1, whereas Al/output signals were observed in the piriform cortex (PIR) and ENT. This input-output correspondence was also conserved in non-mammalian amniotes: in avians, Pr/input signals were enriched in sensory input regions Field L (L), the intercalated hyperpallium (IH), and the entopallium (E), while Al/output signals were enriched in output regions hyperpallium apicale (HA) and nidopallium laterale (NL); in turtles, Pr/input signals localized to the anterior dorsal ventricular ridge (aDVR) and Al/output signals to the medial dorsal pallium (mDP) and lateral pallium (LP). These findings indicate that amniotes share a conserved genetic program that governs the hierarchical organization of cortical sensory circuits. Notably, the avian intermediate mesopallium (MI) region displayed prominent Al/output molecular signatures, contrary to the conventional view that it functions as an association cortex^30^, indicating that this region may serve an output function in some sensory circuits (**Fig. 1E**).

Notably, the same mammalian-derived Pr-Al scoring strategy did not reveal a clear tissue-level Pr-Al gradient in non-amniote species, including axolotl, lungfish, zebrafish, and lamprey (**Fig. 1D-E**). This result may indicate that the mammal-like tissue-level Pr-Al transcriptional gradient was consolidated in amniotes. However, it also reflects an important limitation of this analysis: because the scoring gene sets were defined from mammalian conserved Pr-Al programs, they may preferentially capture molecular features that were retained or amplified in amniotes, while missing more ancient Pr-Al-related features that are conserved at deeper evolutionary levels^10, 35^.

Therefore, to overcome this limitation, we next asked whether more deeply conserved Pr- and Al-like features could be identified at the single-cell level across sarcopterygian vertebrates. This analysis allowed us to determine the deepest evolutionary origin of the Pr-Al axis.

### Pr and Al like cell types conserved in sarcopterygians animals

While spatial transcriptomics revealed an amniote-specific tissue-level gradient, it remained unclear whether Pr-Al-associated molecular signatures exist at the cellular level in more basal vertebrates. To address this, we integrated snRNA-seq datasets of glutamatergic (Glu) and GABAergic (GABA) neurons from six sarcopterygian species spanning over 420 million years of evolution. After standardized data processing and batch correction, we performed consensus cell-type annotation across all species, resolving 7 conserved Glu neuron subclasses and 10 conserved GABA neuron subclasses (**Fig. 2A,B; fig. S3A-F**). Glu subclasses were named based on their laminar distribution and projection identities from mouse and marmoset (**fig. S3B**), including Upper-IT, Middle-IT, Deep-IT, Deep-CT, Deep-PT, Deep-NP, and Deep-RGS12. GABA subclasses were classified according to their developmental origins and canonical marker gene expression, encompassing LGE-derived (LGE.FOXP1, LGE.TSHZ1), CGE-derived (CGE.RELN, CGE.LAMP5-RELN, CGE.LAMP5-PRKG1, CGE.LAMP5, CGE.VIP), and MGE-derived (MGE.SST, MGE.PVALB-POSTN, MGE.PVALB) populations.

**Figure 2.**
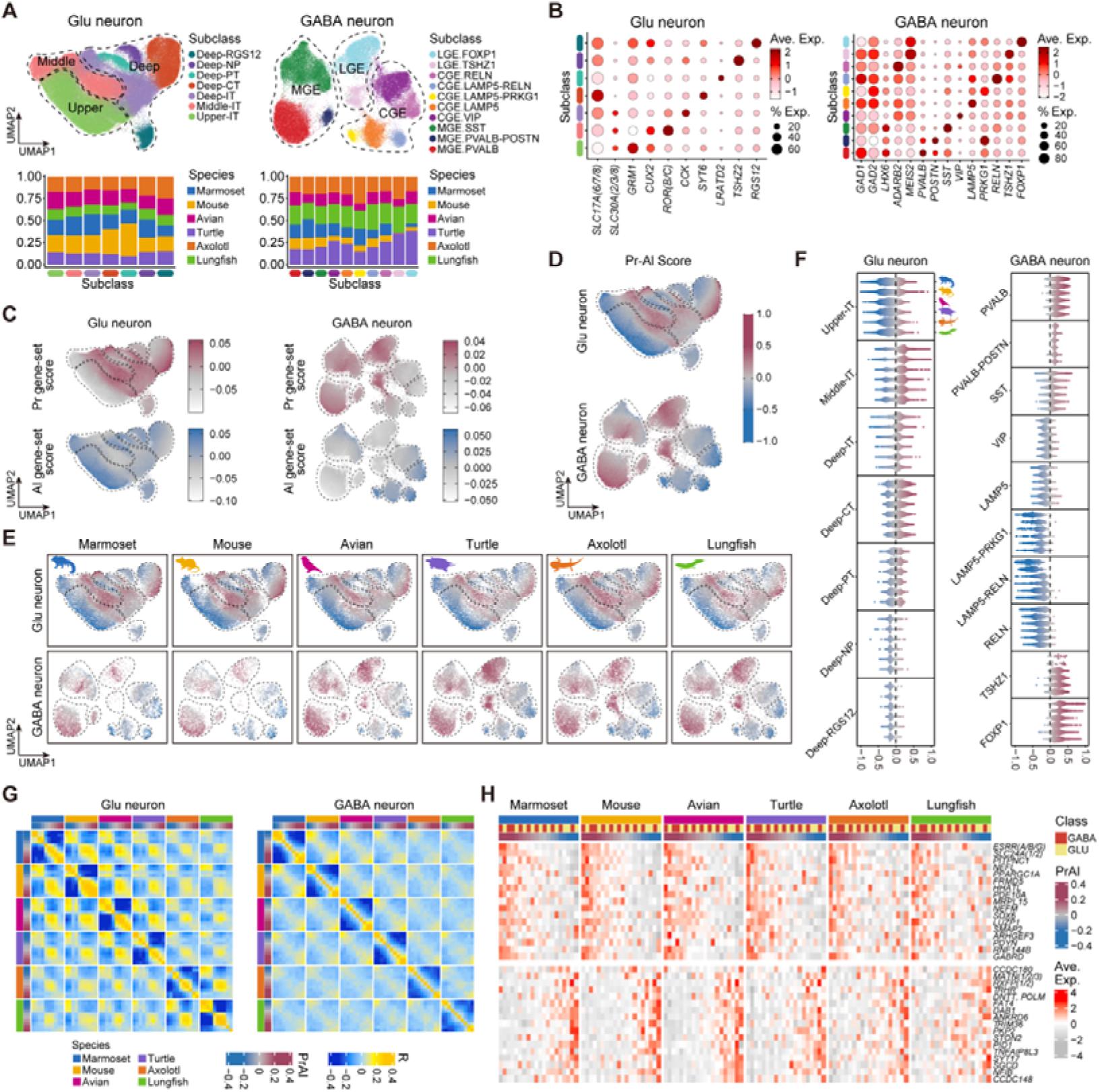
Conserved Pr- and Al-like cell type gradients across sarcopterygians vertebrates. **(A)** UMAP embeddings of integrated glutamatergic (Glu, left) and GABAergic (GABA, right) neuron datasets from six species, colored by cell subclass (top). Bar plots show different subclasses’ ratios in different species (bottom). **(B)** Dot plots showing the average expression and the percentage of cells expressing canonical marker genes for each Glu (left) and GABA (right) subclass. **(C)** Integrated UMAP of Glu (left) and GABA (right) neurons colored by Pr-related gene set score (top) and Al-related gene set score (bottom), demonstrating opposing expression gradients. **(D)** Integrated UMAP of Pr-Al scores in Glu (top) and GABA (bottom) neurons. **(E)** Multi-species UMAP of Glu (top) and GABA (bottom) neurons colored by Pr-Al score, showing conserved gradients across all six species. **(F)** Violin plots of Pr-Al scores across Glu (left) and GABA (right) subclasses in six species. **(G)** Multi-species correlation matrices of cell-type similarity based on Pr-Al score profiles for Glu (left) and GABA (right) neurons. **(H)** Heatmap showing conserved Pr- and Al-related orthologous genes across all six species in scRNA-seq datasets.

Cell type annotations from the cross-species integrated snRNA-seq datasets were robust across all six species. A highly consistent annotation framework was maintained across all taxa, validated by the expression of canonical cell type markers^37,38^ (**Fig. 2B; fig. S3A-F**). Consistent with previous reports^39–41^, we observed relatively fewer LGE-derived GABA neurons in mammals compared with other species (**Fig. 2A; fig. S3A**).

To assess the evolutionary conservation of the Pr-Al index at single-cell resolution, we used the mammalian-conserved Pr and Al orthogroup sets (**Fig. 1C**) to calculate enrichment scores for individual cells in the cross-species integrated snRNA-seq dataset. Notably, in contrast to the spatial transcriptomics results, we observed mutually exclusive, anti-correlated expression patterns of Pr and Al gene sets in both Glu and GABA neurons across all examined species (**Fig. 2C; fig. S3G**). We further quantified this molecular dichotomy using a standardized Pr-Al score, which ranges from +1.0 (pure Pr-like cellular identity) to −1.0 (pure Al-like cellular identity) (**Fig. 2D**).

The Pr-Al score gradient was universally present across all six species, from the basal lungfish to the primate marmoset, demonstrating deep evolutionary conservation of this molecular axis (**Fig. 2E**). Certain cell subclasses showed preferential enrichment at one of the two poles. For example, CGE.RELN interneurons were highly enriched at the Al pole, whereas MGE.PVALB interneurons were concentrated at the Pr pole. Virtually all cell subclasses exhibited a continuous, graded distribution along the Pr-Al axis rather than discrete clustering **(Fig. 2F; fig. S3H-I**). This gradient pattern was independently validated by the expression of spatial Pr- and Al-related genes in single-cell datasets from marmoset and mouse^21^ (**fig. S3J**).

Cross-species correlation analysis of cell types grouped by Pr-Al score revealed that corresponding Pr/Al-related cell types maintained relatively high transcriptomic similarity across species (**Fig. 2G**). Furthermore, we identified a core set of orthologous genes with conserved Pr- or Al-associated expression patterns across all six species. Pr-related genes were predominantly enriched for ion transporters (e.g., *Slc24a1, Slc24a2*) and regulators of neuronal excitability, consistent with the role of primary sensory cortices in rapid signal processing^42^. In contrast, Al-related genes were enriched for factors involved in neuronal migration, laminar patterning, and synaptic plasticity (e.g., *Dab1*), reflecting the specialized functions of the allocortex in complex information processing^43, 44^ (**Fig. 2H**).

Collectively, these results reveal that Pr/Al-related cell-type signatures are not an innovation exclusive to amniotes. Instead, this organizational feature is deeply conserved across sarcopterygian brains, having originated before the divergence of amniotes and amphibians more than 420 million years ago.

This raises a critical question: why do non-amniotes lack a clear Pr-Al transcriptional antagonism in their tissue-level spatial gene expression (**Fig. 1D**)? One plausible hypothesis is that the conserved Pr- and Al-like cell populations identified in these taxa do not exhibit a mutually exclusive spatial distribution aligned with input-output brain regions.

### A conserved Pr-Al index in tetrapod animals

To test this hypothesis and trace the evolutionary origin of the Pr-Al spatial axis, we performed spatial deconvolution analysis across six representative sarcopterygian species. We first divided the integrated Glu and GABA neuron subclasses into ten equal-sized bins based on their Pr-Al scores, generating gradient-resolved cell types ranging from Pr-enriched (e.g., Upper-IT.1, CGE.VIP.1) to Al-enriched (e.g., Upper-IT.10, CGE.VIP.10). We then mapped these gradient-resolved types onto spatial transcriptomic slices of each species using the RCTD algorithm^45^ (**fig. S4A**) and calculated the enrichment score of Pr-Al cell types (**fig. S4B**).

Spatial mapping revealed a striking, evolutionarily conserved pattern across all tetrapod species examined (marmoset, mouse, avian, turtle, and axolotl). Pr- and Al-enriched cell types showed significantly anti-correlated spatial distributions (R = −0.34 to −0.71) (**Fig. 3A, fig. S4C**), and selectively localized to cortical input and output regions, respectively (**Fig. 3B-C**). Remarkably, the axolotl — previously thought to lack a mammal-shared Pr-Al gradient based on spatial transcriptomic data — also exhibits this conserved organizational pattern, with Pr cell types enriched in the ventral pallium (VP) and Al cell types enriched in the dorsal pallium (DP).

**Figure 3.**
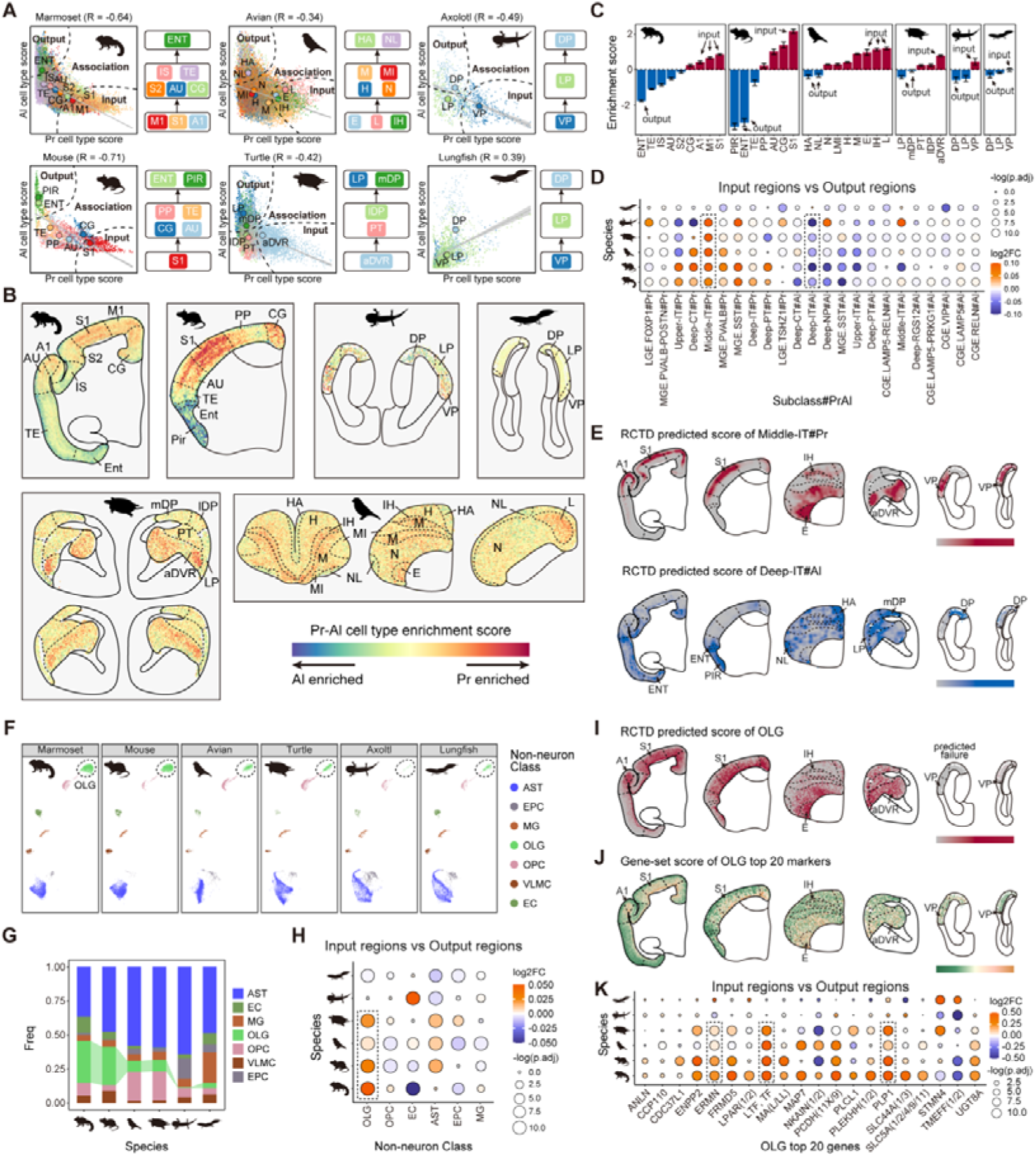
Conserved Pr-Al spatial axis in tetrapods and its evolutionary origin. **(A)** Violin plots of Pr-Al scores for integrated glutamatergic (left) and GABAergic (right) neurons, grouped into ten equal-sized bins. Schematic illustrating the RCTD spatial deconvolution approach for mapping gradient-resolved cell types onto brain slices. **(B)** Spatial distribution of Pr-Al cell type enrichment scores across six sarcopterygian species. **(C)** Enrichment scores of Pr (red) and Al (blue) gene sets in cortical input and output regions across species. Major input and output brain regions are annotated for each species. **(D)** Dot plot showing log2 fold change (log2FC) and significance (-log10 adjusted p-value) of Pr-Al gradient cell types between input and output regions across species. **(E)** RCTD-predicted spatial scores of representative conserved cell types: Pr-related Middle-IT (upper) and Al-related Deep-IT (lower). See **fig. S4D** for other cell type. **(F)** Scatter plots of annotation for non-neuronal cell types across six species. OLG, oligodendrocyte. AST, astrocyte; EC, endothelial cell; MG, microglia; OPC, oligodendrocyte precursor cell; VLMC, vascular leptomeningeal cell. **(G)** Bar plot shows that frequency distribution of non-neuronal cell types in each species. **(H)** Dot plot showing log2FC and significance of non-neuronal cell types between input and output regions across species. **(I)** RCTD-predicted spatial scores of OLGs across species. **(J)** Gene set scores of the top 20 OLG marker genes across species. **(K)** Dot plot showing log2FC and significance of the top 20 OLG marker genes between input and output regions across species.

In stark contrast, lungfish—the closest living relatives of tetrapods—showed no spatial segregation of Pr- and Al-enriched cell types, nor any anti-correlation between Pr-Al scores (R = 0.39) (**Fig. 3A-C; fig. S4C**). Quantitative analysis of cell type enrichment between input and output regions confirmed these observations. We identified a panel of Pr- and Al-related cell types uniformly enriched in their respective brain regions across all tetrapods (**Fig. 3D**). For example, Middle-IT#Pr and Deep-IT#Al cells, which mediate subcortical input and output in mammals^21, 46^, were universally enriched in Pr input and Al output regions, respectively. However, no such conserved enrichment pattern was detected in lungfish (**Fig. 3D-E**).

Intriguingly, we also identified several lineage-specific Pr- or Al-associated neuron subtypes, including primate-specific (Upper-IT#Pr, Deep-PT#Pr), amniote-specific (Deep-NP#Al, PVALB#Pr), and tetrapod-specific (Deep-IT#Pr, MGE.SST#Pr, Upper-IT#Al, Deep-PT#Al) subtypes, whose evolutionary origins and functional roles in neural circuits warrant further investigation (**fig. S4D**).

Collectively, these results demonstrate that although lungfish possess Pr-Al-associated cell types at the single-cell gene expression level, they have not established a spatially organized Pr-Al axis. This indicates that the Pr-Al spatial axis arose after the water-to-land transition, coinciding with the origin of tetrapods approximately 390 million years ago.

### Amniote-specific recruitment of Pr-related oligodendrocytes

Having established the evolutionary origin of the neuronal Pr-Al spatial axis in tetrapods, we next investigated whether non-neuronal cell types also contribute to the functional specialization of this axis. Given previous reports that oligodendrocytes (OLGs), the myelinating cells of the central nervous system, are preferentially enriched in Pr regions to support rapid signal conduction in sensory areas^47^, we examined the spatial distribution of non-neuronal cell types across species.

Integrated analysis of non-neuronal cells revealed a progressive increase in OLG proportions along the sarcopterygian lineage: marmoset and mouse showed the highest OLG proportions, followed by avian and turtle, whereas axolotl and lungfish had markedly reduced OLG proportions (**Fig. 3F-G**). Spatial enrichment analysis showed that OLGs were selectively and consistently enriched in Pr-dominant input regions across all amniote species (marmoset, mouse, avian, turtle), but not in axolotl or lungfish (**Fig. 3H-I**). Furthermore, the top 20 OLG marker genes, including the canonical myelin genes *PLP1* and *ERMN*, also showed significant enrichment in Pr regions exclusively in amniotes^38, 48^ (**Fig. 3J-K**). This is consistent with prior experimental findings: the axolotl pallium lacks robust myelination^49^.

These findings suggest that the recruitment of OLGs to Pr regions, which likely enhanced the functional specialization of the Pr-Al input-output axis by accelerating sensory signal conduction, arose later in amniote evolution.

### Evolutionary conservation and stepwise assembly of the Pr-Al molecular axis across tetrapods

Having established that a spatially organized Pr-Al molecular axis, corresponding to cortical input-output hierarchies, is conserved across tetrapods but absent in non-tetrapod vertebrates, we first performed Mfuzz clustering^50^ of conserved orthologs across five tetrapod species to define its molecular basis and evolutionary origins. We identified Pr-enriched (orange) and Al-enriched (blue) gene clusters (**Fig. 4A, fig. S5A**). This analysis revealed both deeply conserved and species-specific expression patterns. For example, consistent with previous reports, the axon guidance ligand *SEMA7A* and its receptor *PLXNC1* exhibited mutually exclusive, anti-correlated expression along the Pr-Al axis in mouse and avian^22^, and were confirmed as Pr-enriched and Al-enriched genes, respectively, in our dataset. In contrast, the transcription factor *ZBTB40* showed Pr-enrichment only in mouse, whereas the cadherin *CDHR1* was Al-enriched in mouse but Pr-enriched in avian, demonstrating lineage-specific functional divergence^51, 52^ (**Fig. 4A**).

**Figure 4.**
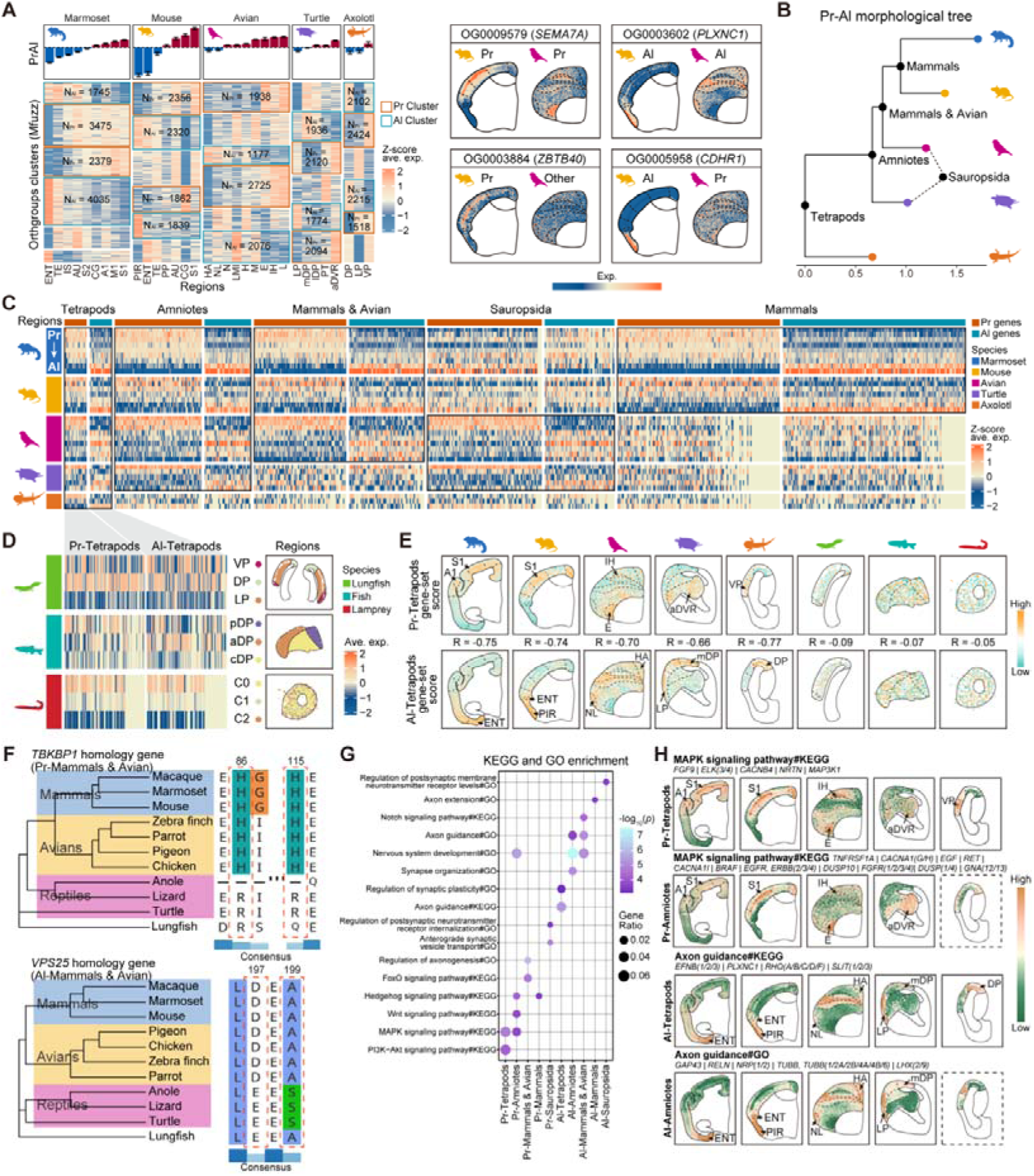
Evolutionary conservation and molecular basis of the Pr-Al axis. **(A)** Mfuzz clustering of conserved orthologs into Pr-enriched (orange) and Al-enriched (blue) clusters across five tetrapod species. Brain regions are ordered from Al-dominant to Pr-dominant along the x-axis. Right panels show spatial expression patterns of representative genes. **(B)** Tree constructed from Pr-Al gene sets across species using the maximum likelihood method. For alternative tree construction methods, see **fig. S5B**. **(C)** Heatmap showing the expression of evolutionarily stratified Pr-Al gene sets across brain regions of all tetrapod species. Gene sets include tetrapod-conserved, amniote-conserved, mammals-avian conserved, sauropsid-specific, and mammals-specific. **(D)** Heatmap show the regional distribution of tetrapod-conserved Pr and Al gene sets (from C) in lungfish, zebrafish, and lamprey. **(E)** Spatial distribution of tetrapod-conserved Pr-related (top) and Al-related (bottom) gene set scores (from C) across all vertebrate species. Pearson correlation coefficients (R) between the two gene sets are indicated in the middle. **(F)** Protein sequence alignments of *TBKBP1* (top, Pr-Mammals & Avian) and *VPS25* (bottom, Al-Mammals & Avian), showing convergent amino acid substitutions specific to mammals and avian: H86R and H115R in *TBKBP1*, and D197E and A199S in *VPS25*. **(G)** GO and KEGG enrichment analysis of evolutionarily stratified Pr-Al gene sets. **(H)** Spatial distribution reveals the scores of representative biological-function gene sets in the enrichment analysis (from G) corresponding to certain representative genes across all tetrapod species.

Strikingly, a phylogenetic tree constructed from the Pr-Al gene set did not recapitulate the canonical tetrapod phylogeny (**Fig. 1B**). Instead, mammals and avians formed a distinct clade separate from reptiles and amphibians (**Fig. 4B, fig. S5B**). This unexpected topology indicates that mammals and avians may share more similar Pr-Al molecular signatures than expected from their phylogenetic distance, providing some evidence for convergent molecular evolution of this brain organizational feature in the two amniote lineages. This observation prompted us to systematically define five evolutionarily stratified Pr-Al gene sets: tetrapod-conserved, amniote-conserved, mammals-avian conserved, sauropsid-specific, and mammal-specific (**Fig. 4C**). Spatial transcriptomic (**fig. S5C**) and snRNA-seq (**fig. S6A**) analyses validated the regional and cellular enrichment, respectively, of all evolutionarily stratified Pr-Al gene sets across vertebrate species.

Heatmap analysis revealed no mutually exclusive, anti-correlated regional expression between tetrapod-conserved Pr and Al gene sets in lungfish, zebrafish, and lamprey (**Fig. 4D**). Consistently, spatial scoring of these gene sets failed to detect a robust Pr-Al molecular gradient, with Pearson correlation coefficients ranging from −0.09 to −0.05 (**Fig. 4E**). Together, these findings suggest that the Pr-Al molecular axis represent a tetrapod-specific innovation that arose following the water-to-land transition.

To identify sequence-level changes underlying the observed convergent molecular evolution, we performed comparative genomic analysis specifically on the mammals-avian conserved Pr-Al gene set (**Fig. 4F, fig. S6B**). This analysis identified convergent amino acid substitutions in multiple genes, including the Pr-associated *TBKBP1* (H86R and H115R), a regulator of synaptic vesicle trafficking and neuronal innate immune responses^53^, and the Al-associated *VPS25* (D197E and A199S), a core component of the ESCRT-II complex that mediates synaptic glutamate receptor recycling and endosomal sorting^54^ (**Fig. 4F**). Additional representative convergent substitutions were found in *DBX2* (Pr-mammals & avian), a homeodomain transcription factor that controls glutamatergic neuron fate specification during cortical development^55^; *TIMM29* (Al-Mammals & Avian), a mitochondrial inner membrane translocase subunit essential for high-energy-demand neuronal function and mitochondrial homeostasis^56^; and *RELT* (Pr-Mammals & Avian), a member of the tumor necrosis factor receptor superfamily that plays a vital modulatory role in the normal assembly of central neural circuits^57^ (**fig. S6B**). Notably, these convergent substitutions were concentrated in genes with essential functions in core neuronal processes, indicating that parallel protein sequence changes have broadly shaped the convergent evolution of the Pr-Al molecular axis in mammals and avians.

Functional enrichment analysis revealed the stepwise assembly of the Pr-Al molecular program (**Fig. 4G-H, fig. S6C**). The MAPK signaling pathway, which regulates sensory system functions^58–61^, was strongly enriched in both tetrapod-conserved and amniote-conserved Pr gene sets. The tetrapod core module included *FGF9* and *NRTN*, critical for neurogenesis and peripheral nervous system development, supporting terrestrial locomotion and digestion^62, 63^, as well as *CACNB4*, which mediates presynaptic calcium influx and neurotransmitter release^64^. The amniote module further expanded with additional regulators of synaptic plasticity and neurotransmitter release, such as *CACNA1* and *CACNA1G*^64^.

Axon guidance pathways were enriched across all evolutionary strata, with progressive expansion: tetrapod-conserved genes included *EFNB1* and *PLXNC1*, which establish topographic projections^65–67^; amniote-conserved genes added *GAP43* and *RELN*, which regulate cortical lamination^43, 68, 69^; and mammals-avian conserved genes included *BDNF* and *NTF3*, two canonical neurotrophins that support neuronal survival, neurite extension, and circuit maturation^70, 71^. Lineage-specific enrichments included the FoxO signaling pathway, which includes *ATM* and *TGFBR2*, both involved in neuronal stress resistance in mammals and avians^72, 73^. The Hedgehog pathway, which includes *USP9X* and *HSP90AA1*, suggests additional developmental signaling and proteostasis mechanisms that may support synaptic maturation^74, 75^.

Collectively, these results demonstrate that the Pr-Al index is a deeply conserved organizational feature of the tetrapod brain. Its core molecular module, centered on MAPK signaling and axon guidance, originated in the last common tetrapod ancestor and was subsequently expanded and refined during amniote and mammalian evolution to support increasing neural circuit complexity and cognitive function.

### Convergent molecular signatures of association cortices in mammals and avians

We next examined the correlation between the Pr-Al index and the expansion of association cortices, which support advanced cognitive functions^21, 23^. We first generated whole-pallium flatmaps of marmoset and mouse to visualize the Pr-Al index and the derived intersection index, a metric previously linked to association regions^21^. We then defined Mammalian Association Genes (MAGs): genes conserved between marmoset and mouse and specifically enriched in association cortices. Spatial scoring confirmed MAGs were highly enriched in prefrontal, parietal, and temporal association regions of both species, precisely overlapping with high intersection index areas (**Fig. 5A**).

**Figure 5.**
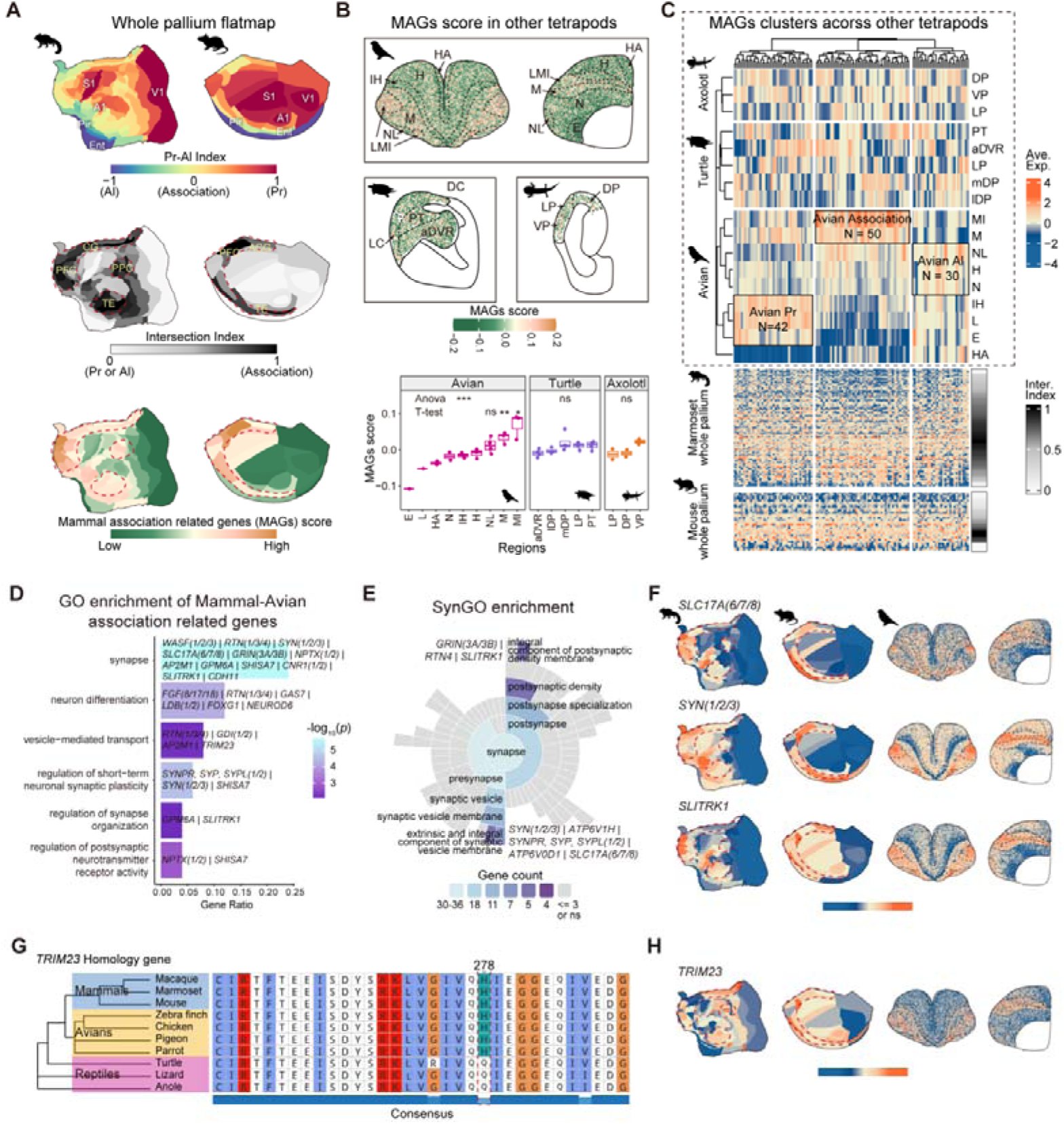
Convergent evolution of association cortex molecular signatures in mammals and avians. **(A)** Whole-pallium flatmaps of common marmoset and mouse showing the Pr-Al index, intersection index, and spatial scores of mammals association-related genes (MAGs). **(B)** Top, spatial distribution maps of MAG scores in avian, turtle, and axolotl. Bottom, box plots showing statistical comparisons of MAG scores across brain regions using one-way ANOVA followed by post hoc t-tests. *, p < 0.05; **, p < 0.01; ***, p < 0.001; ns, not significant. **(C)** Heatmap of MAG expression across tetrapod brain regions, revealing convergent association cortex molecular clusters that are shared by mammals and avians. **(D)** Gene Ontology (GO) enrichment analysis of the convergent mammals-avian association-related gene set. **(E)** Synaptic Gene Ontology (SynGO) enrichment analysis showing significant enrichment of the convergent gene set in presynaptic and postsynaptic components. **(F)** Spatial expression patterns of representative synaptic genes including *SLC17A6/7/8*, *SYN1/2/3*, and *SLITRK1* in marmoset, mouse, and avian. **(G)** Protein sequence alignment of *TRIM23* showing the convergent H278Q amino acid substitution that is specific to mammals and avian. **(H)** Spatial expression patterns of *TRIM23* across species, confirming its preferential enrichment in association cortices.

To test whether this molecular signature exists in other tetrapods, we scored the MAGs across spatial transcriptomic datasets from avian, turtle, and axolotl. Strikingly, only avians showed significant regional enrichment (ANOVA, *p* < 0.001), with highest scores in the M and MI—regions functionally analogous to mammalian association cortices^30, 76^. Statistical analysis confirmed that MAG scores in M and MI were significantly higher than in all other avian brain regions, whereas no such enrichment was observed in turtle or axolotl (**Fig. 5B, fig. S7A**). Unsupervised clustering of MAG expression across all tetrapods revealed a distinct gene cluster consistently enriched in association regions of mammals and avians, but not in other tetrapods (**Fig. 5C**).

Functional enrichment analysis of this convergent gene set showed significant enrichment of genes involved in neuronal function, particularly synapse organization, synaptic transmission, and neuronal differentiation (**Fig. 5D, fig. S7B**). SynGO enrichment^77^ further distinguished presynaptic and postsynaptic components: presynaptic genes included *SLC17A6/7/8*, responsible for glutamate loading into synaptic vesicles^78^, and *SYN1/2/3*, that regulate synaptic vesicle release^79^; postsynaptic genes included *SLITRK1*, crucial for synapse formation and plasticity^80^ (**Fig. 5E**). Spatial visualization confirmed the conserved enrichment of these representative genes in the association cortices of marmoset, mouse, and avian (**Fig. 5F, fig. S7C**). These findings suggest that the convergent evolution of advanced cognitive abilities in mammals and avian was supported by the independent deployment of a conserved molecular module for synaptic function and plasticity.

To identify sequence-level changes underlying this convergence, we performed comparative genomic analysis of the convergent MAG set. *TRIM23*, a vesicle-trafficking factor associated with Golgi and lysosomal transport, may contribute to neuronal developmental programs underlying this convergence^81, 82^, exhibited a notable convergent amino acid substitution at position 278: all mammals and avian shared a histidine (H), while all reptiles and amphibians had a glutamine (Q) (**Fig. 5G**). Spatial expression confirmed *TRIM23* was highly enriched in the association cortices of mammals and avian (**Fig. 5H**), suggesting this mutation may have enhanced synaptic function in these lineages.

Collectively, our results demonstrate that mammals and avian independently evolved similar molecular signatures in their association cortices, centered on genes regulating synaptic transmission and plasticity. This convergent molecular evolution provides a mechanistic basis for the parallel emergence of advanced cognitive abilities in amniotes.

## Discussion

Our study provides a cross-species molecular framework for understanding the evolution of pallial circuitry. By integrating spatial transcriptomic and single-nucleus transcriptomic data across major vertebrate lineages, we show that the Pr-Al axis emerged through a stepwise evolutionary process. Pr- and Al-like cellular identities are deeply conserved across sarcopterygians, but their spatial segregation into an input-output axis appears only in tetrapods. A robust tissue-level Pr-Al transcriptional gradient subsequently became a shared feature of amniote pallia, where it aligns with sensory-processing hierarchies across both the mammalian cortex and the sauropsid pallium ^21^.

These findings help resolve a long-standing paradox in comparative neurobiology. Although the DVR and neocortex are not structurally homologous, they share striking similarities in sensory circuit organization^7, 13, 14^. Our results suggest that this paradox can be reconciled by separating structural origin from molecular circuit logic. The conserved Pr-Al program does not imply that the DVR and neocortex arise from the same pallial territory; rather, it indicates that independently derived pallial regions can deploy homologous cell-type and gene-regulatory programs to construct analogous input-output architectures. In this sense, the Pr-Al axis provides a quantitative molecular bridge between classical anatomical models and modern cell-type homology^19^.

The evolutionary timing of the Pr-Al axis further suggests that the water-to-land transition marked a major inflection point in vertebrate brain organization. Lungfish retain Pr- and Al-like neuronal states at single-cell resolution, but lack their spatial alignment with input-output regions. Tetrapods, by contrast, organize these conserved cell states into a coherent regional axis. This observation implies that the origin of terrestrial vertebrates involved not only morphological and sensory adaptation, but also the spatial redeployment of ancestral neuronal programs into ordered circuit maps. The later amniote-specific enrichment of oligodendrocytes in Pr regions may have further sharpened this axis by promoting faster sensory information transfer through increased myelination^47^.

Our evolutionary stratification of Pr-Al-associated genes further suggests that this axis was assembled through a combination of deep conservation and lineage-specific innovation. A tetrapod core enriched for MAPK signaling and axon guidance may have provided the initial molecular scaffold for regionalized input-output organization, whereas amniote and mammal-bird modules added genes involved in synaptic regulation, circuit refinement, myelination, and proteostasis. The unexpected similarity between mammalian and avian Pr-Al molecular programs, together with convergent protein-sequence changes, indicates that parallel pallial evolution operated not only at the level of gross anatomy, but also at the levels of gene expression and protein evolution.

The convergent molecular signature of mammalian and avian association regions has important implications for the evolution of cognition. Birds lack a laminated neocortex, yet they exhibit sophisticated cognitive abilities supported by nuclear pallial architectures^30, 76^. Our data show that avian association-like pallial regions are enriched for a mammalian association-gene program dominated by synapse organization and neurotransmission. This observation suggests that advanced pallial computation may depend less on a specific laminar scaffold than on the deployment of conserved synaptic and cell-type programs within expanded associative networks.

Several limitations should be noted. First, cross-species transcriptomic comparisons depend on orthogroup definitions and cannot fully resolve lineage-specific paralogs or regulatory innovations. Second, differences in spatial transcriptomic resolution and anatomical registration across species may obscure finer subregional organization. Third, our current analyses remain restricted to sampled pallial regions and therefore lack validation at the whole-brain level across species. Future studies incorporating whole-brain spatial and single-cell datasets will be important for testing whether the Pr-Al framework extends beyond the currently sampled territories and for assessing its relationship to broader vertebrate brain organization. Fourth, the convergent sequence changes identified here remain candidate mechanisms and require direct functional validation. More generally, future work integrating developmental time courses, chromatin accessibility, projection mapping, and perturbation experiments will be essential for determining how Pr-Al gene modules regulate circuit assembly and whether mammal-bird convergent substitutions alter synaptic or regional specialization. Nevertheless, our results establish the Pr-Al axis as an evolutionarily informative framework for understanding how conserved molecular programs can be redeployed to generate parallel pallial circuit architectures.

## Materials and Methods

### Collection and preprocessing of spatial transcriptomic data

To perform a comparative analysis of the Pr-Al index across vertebrates spanning 500 million years of evolution, we collected published Stereo-seq spatial transcriptomic (ST) data of the adult pallium from 8 species: 3 sagittal sections for sea lamprey, 2 sagittal sections for zebrafish, 1 coronal section covering both hemispheres for West African lungfish, 1 coronal section covering both hemispheres for Mexican axolotl, 2 coronal sections covering both hemispheres for Chinese softshell turtle, 1 bilateral hemisphere section plus 2 hemicoronal sections for zebra finch, 1 representative hemicoronal section for mouse, and 1 representative hemicoronal section for common marmoset.All sections were processed using the Stereo-seq Analysis Workflow (SAW). All SAW processing steps were executed via a wrapped pipeline on the DCS cloud platform (https://cloud.stomics.tech/)^83, 84^.

Based on the original anatomical annotations of each dataset, we manually excluded the hippocampus and its homologous structure, the medial pallium (MP; in turtle, axolotl and lungfish), as well as the ventral pallium (VP) in fish^10^. The remaining pallial regions were retained for downstream analyses. Unsupervised clustering was performed separately for each species using the Seurat v4.4.0 package^85^. For species with multiple sections (3 for avian, 2 for turtle), additional batch correction across sections was conducted using the Harmony v1.2.1 package^86^.

Clustering parameters were adjusted individually for each species to match the actual distribution of anatomical brain regions, including the number of highly variable genes (3000 for avian, 5000 for turtle, 2000 for axolotl, 1500 for lungfish, 3000 for zebrafish, 3000 for lamprey), the number of principal component analysis (PCA) dimensions (10 for avian, 10 for turtle, 10 for axolotl, 15 for lungfish, 15 for zebrafish, 15 for lamprey), and clustering resolution (0.8 for avian, 0.2 for turtle, 0.2 for axolotl, 0.8 for lungfish, 0.5 for zebrafish, 0.5 for lamprey). Combined with clustering results, manual anatomical annotation of brain regions was performed using ImageJ software^87^.

Notably, some avian anatomical regions showed highly similar expression profiles in clustering and could not be directly distinguished, including nidopallium (N) vs. hyperpallium (H), field L (L) vs. intercalated hyperpallium (IH) vs. entopallium (E), and hyperpallium apicale (HA) vs. nidopallium laterale (NL). We therefore performed manual anatomical annotation based on the topological positions of each region: N and NL lie ventral to region M, whereas H and HA lie dorsal to region M.

### Identification of orthogroups

To establish a unified framework for cross-species gene expression analysis, we first collected coding sequence (CDS)-derived protein sequences from the reference genome of each species in National Center for Biotechnology Information (NCBI, https://www.ncbi.nlm.nih.gov/): marmoset (GCF_009663435.1), mouse (GCF_000001635.27), zebra finch (GCF_048771995.1), Chinese softshell turtle (GCF_000230535.1), axolotl (AmexG_v6.0-DD), West African lungfish (GCF_019279795.1), zebrafish (GCF_049306965.1), and sea lamprey (GCA_015708825.1) . The longest transcript per gene was selected as the representative sequence.

Orthogroup identification was then performed using OrthoFinder v2.5.5^34^, yielding a total of 37,487 orthogroups. Detailed statistics are provided in **Fig. S2A-B**. For each orthogroup, expression counts of its member genes in the ST and single-nucleus RNA-seq (snRNA-seq) matrices were summed to generate a spot/cell × orthogroup expression matrix.

### Scoring and cross-Species comparison of Pr and Al gene sets in spatial transcriptomic data

To identify conserved orthogroups in mammals for unified Pr-Al scoring across all species, we obtained published region × gene whole-brain expression matrices of mouse and marmoset^21^, and constructed region × orthogroup matrices via orthogroup mapping.

Using the Pr-Al index defined in the original study, we calculated the Pearson correlation coefficient between each orthogroup and the Pr-Al index. Orthogroups with Pearson correlation coefficient *R* > 0.5 and *R* < −0.5 were defined as Pr-associated and Al-associated high-confidence orthogroups, respectively, yielding conserved Pr and Al orthogroup sets shared between mouse and marmoset.

Based on these two orthogroup sets, gene set enrichment scoring was performed on log-normalized and Z-score transformed ST matrices of each species using the Seurat “AddModuleScore” function.

### Definition and classification of sensory hierarchical levels across species

To enable unified cross-species comparison of sensory hierarchy, we integrated published experimental evidence and review articles for each species, and classified pallial regions into three hierarchical tiers — input, association and output — for comparative analysis.

1. Mammals: The sensory circuitry of the pallium has been well defined in the neocortex, allocortex and periallocortex. Primary sensory cortices serve as the core input regions, including the primary somatosensory/motor cortex (S1/M1), primary auditory cortex (A1) and primary visual cortex (V1). Regions such as the prefrontal, temporal and posterior parietal cortices act as association cortices, participating in secondary processing of sensory signals^46, 88, 89^. The entorhinal cortex (ENT) and piriform cortex (PIR) receive integrated multimodal sensory signals^90–92^, and function as output regions projecting to the hippocampus and other subcortical nuclei.
2. Reptiles: Although the anterior dorsal ventricular ridge (aDVR) is considered homologous to the mammalian lateral amygdala and claustrum, it functions similarly to mammalian sensory input regions and receives extensive thalamic sensory input^93–95^. Previous work has characterized the visual hierarchy in turtles^96^, the medial dorsal pallium (mDP) acts as an output region, relaying visual signals to the hippocampus; the D1 and D2 regions (corresponding to the lateral dorsal pallium, lDP, and pallial thickening, PT, respectively^31^) occupy an intermediate position and likely serve association functions. The lateral pallium (LP) has not been systematically characterized in terms of circuit position, but given its partial homology to the mammalian PIR^9, 97^, we classified it as an output region, consistent with the status of the mammalian PIR.
3. Avian: Regions E, L and IH are recognized as visual, auditory and multimodal sensory input regions, respectively^30, 98^. Retrograde tracing studies have confirmed that HA^99^ and NL (including the nidopallium caudolaterale, NCL; nidopallium frontolaterale, NFL; and nidopallium intermedium laterale, NIL^100^) may lie at the end of the sensory circuit, analogous to output regions. The remaining nidopallium, hyperpallium and entire mesopallium are generally classified as association regions^98^.
4. Amphibians: The medial pallium (MP), the homologue of the hippocampus, both receives extensive sensory input and mediates sensory output from the pallium^32^; this feature may represent an evolutionary innovation of the amphibian thalamo-pallial circuit. The ventral pallium (VP) receives sensory projections from the MP as well as from the anterior and central thalamic nuclei^32^, and is functionally more analogous to a canonical input region in terms of circuit architecture. Regarding connectivity patterns, the lateral pallium (LP) connects to both the VP and dorsal pallium (DP), whereas the VP and DP have no direct connections^101, 102^. Accordingly, in the amphibian pallial sensory hierarchy, the LP and DP serve association-like and output-like functions, respectively.
5. Lungfish: The regional organization and connectivity pattern of the lungfish pallium are highly similar to those of amphibians^33^, so we adopted the amphibian classification scheme for its sensory hierarchy.
6. Ray-finned fish: A recent study identified a sensory hierarchy in the teleost pallium, with a posterior-to-anterior gradient in the dorsal pallium (DP)^103^. However, the primary sensory relay structure upstream of the teleost pallium is not the thalamus, but the preglomerular nucleus (PG), which receives major sensory input and relays signals to the DP^104^.
7. Lamprey: A sensory hierarchy may also exist in the lamprey pallium. Previous work has indicated that visual and somatosensory inputs are mainly distributed in the dorsal lateral pallium^35^, but no systematic hierarchical classification has been established.

### Collection and preprocessing of single-nucleus transcriptomic data

For systematic cross-species comparison of single-cell profiles, we prioritized snRNA-seq datasets paired with the ST studies of each species. These datasets were generated using similar experimental protocols and the same reference genomes, enabling better matching with corresponding ST data. No paired snRNA-seq dataset was available for axolotl, so a published conspecific dataset was used as a substitute^97^.

Prior to unsupervised clustering, snRNA-seq data from each species were converted into orthogroup expression matrices. To equalize sequencing depth across species, each dataset was downsampled to 40,000 cells and deleted a spot of unknown glial cells from lungfish and mix feature of Glutamatergic (Glu) and GABAergic (GABA) neuron after clustering. All downstream analyses were performed using the Seurat package: 5000 anchor genes were first selected, and initial cross-species batch correction was performed via the RPCA workflow of Seurat’s “IntegrateData”; secondary batch correction for all internal batches was then performed using the Harmony algorithm. Finally, two-dimensional UMAP embeddings were generated using the first 30 Harmony dimensions for coarse clustering, yielding 12 major cell classes.

Glu and GABA neurons were then isolated separately for subclustering. Given that interspecies expression differences may exceed differences between cell subtypes, only genes detected in at least 5 species were retained for this step. Gene expression counts were normalized using Seurat’s “SCTransform”, initial interspecies batch effects were removed via “IntegrateData”, and secondary batch correction for internal batches was again performed with Harmony.

Both datasets were subjected to unsupervised clustering using the first 15 Harmony dimensions and a resolution of 0.5. Guided by original cell annotations from mouse and marmoset (**Fig. S3B**), Glu neurons were classified into 7 subclasses, and GABA neurons into 10 subclasses.

### Identification and scoring of the Pr-Al axis in single-cell data

For single-cell analysis, we took the intersection of the top 1000 Pr- and Al-associated genes from marmoset and mouse as the feature gene set, and scored Glu and GABA neurons separately using Seurat’s “AddModuleScore” function on the integrated expression matrix.

To mitigate scoring fluctuations caused by the sparsity of single-cell data, we smoothed the Pr and Al scores separately using a gene imputation workflow adapted from the MAGIC algorithm^105^. Pr and Al scores were then normalized to the 0-1 range within each species, and the single-cell Pr-Al score was calculated according to the Pr-Al index formula:

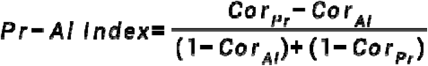

To facilitate cross-species similarity comparison of Pr-Al-associated cells and identification of related genes, Glu and GABA neurons were each divided into 10 equal-sized bins based on their Pr-Al scores. Average expression levels and average Pr-Al scores of the 10 bins were calculated for each species to generate a pseudo-bulk expression dataset. Based on this pseudo-bulk data, we computed Pearson correlation coefficients of Pr-Al-associated cell profiles between pairwise species (**Fig. 2G**), and screened for cross-species conserved single-cell Pr-associated (*R* > 0.25) and Al-associated (*R* < −0.25) genes based on the correlation between gene expression and Pr-Al score (**Fig. 2H**).

### Spatial enrichment analysis of Pr and Al cell types

Based on the 10 equal-sized bins of Glu and GABA neurons, we combined cell subclass identities with bin labels to define gradient-resolved cell types with Pr-Al characteristics (**Fig. S4A**). The spatial distribution of these Pr-Al gradient cell types in ST data was then predicted using the RCTD v2.2.1 deconvolution algorithm^45^, with the doublet_mode parameter set to full.

After deconvolution, the proportion of each cell type per spot was calculated by dividing the predicted score of that cell type by the total score of the spot. The Pr-Al enrichment score for each species was computed as the weighted sum of Z-score normalized cell proportions, with the Pr-Al score of each cell type as the weight. We further defined Pr-type cells (Pr-Al score > 0.1) and Al-type cells (Pr-Al score < −0.1), and calculated the Pr cell type score and Al cell type score separately using the same weighted approach (**Fig. S4B**).

To verify the mutually exclusive enrichment pattern of Pr and Al subtypes of each cell subclass between input and output regions across species, we aggregated RCTD predicted scores for Pr and Al cells of each subclass, and calculated log fold change (log FC) and *P* values between input and output regions using Seurat’s “FindMarkers” function.

For non-neuronal cells, we mixed them with neurons and predicted their spatial distribution via RCTD simultaneously, and also used FindMarkers to compute expression differences between input and output regions. For oligodendrocytes (OLGs), we selected the top 20 marker genes identified from snRNA-seq data, performed gene set scoring with Seurat’s “AddModuleScore”, and quantified their enrichment between input and output regions using “FindMarkers”.

### Identification and evolutionary analysis of Pr- and Al-associated genes in tetrapods

Given the large interspecies differences in pallial size and number of ST spots, we identified Pr- and Al-associated genes at the regional level to balance analytical scale across species. Before constructing regional pseudo-bulk data, we performed gene expression imputation for each species using the MAGIC algorithm to improve detection sensitivity for low-abundance, sparsely expressed genes.

Regions of each species were sorted by Pr-Al enrichment score, and time-series expression clustering was performed using the Mfuzz package^50^. The number of clusters was uniformly set to 6 per species. Based on the degree of differential expression between input and output regions (log FC) and the Pearson correlation between gene expression and regional order (ranked 1 to *n*), we selected 2 Pr-enriched clusters and 2 Al-enriched clusters for each species (**Fig. S5A**).

To construct phylogenetic trees for Pr-Al-associated genes, we categorized gene expression traits into three states: Pr-like, Al-like and other. Three tree-building methods were applied: maximum likelihood tree construction using the “pml_bb” function of the phytools package^106^; maximum parsimony tree construction using the “pratchet” function of phytools; and Bayesian inference using MrBayes^107^.

To validate the five evolutionarily stratified Pr-Al gene sets, we scored them in ST and snRNA-seq data of each species using Seurat’s “AddModuleScore”, to confirm their specific enrichment in input/output brain regions and Pr/Al cells of the corresponding lineages.

### Convergent evolution sequence analysis in mammals and avian

To investigate the phylogenetic relationships of genes across mammals, avian and reptiles, we collected protein sequences from 11 vertebrate species: rhesus macaque (*Macaca mulatta*), common marmoset (*Callithrix jacchus*), house mouse (*Mus musculus*), zebra finch (*Taeniopygia guttata*), chicken (*Gallus gallus*), rock dove (*Columba livia*), budgerigar (*Melopsittacus undulatus*), Chinese softshell turtle (*Pelodiscus sinensis*), common wall lizard (*Podarcis muralis*), green anole (*Anolis carolinensis*) and West African lungfish (*Protopterus annectens*). Lungfish was selected as the outgroup for subsequent phylogenetic analyses.

First, gene names in protein FASTA files of each species were uniformly parsed, and the longest protein sequence per target gene in each species was extracted as the representative sequence. Multiple sequence alignment was then performed for the 11 representative sequences of each shared gene using MAFFT^108^, with parameters set to –localpair --maxiterate 1000. To reduce the impact of low-confidence alignment regions on tree topology, alignments were trimmed using ClipKIT^109^ with the smart-gap trimming mode.

Phylogenetic trees were constructed using IQ-TREE2^110^. Trimmed protein alignments were used as input, the best-fit substitution model was selected via the MFP strategy, and branch support was evaluated using ultrafast bootstrap (-B 1000) and SH-aLRT test (-alrt 1000). The lungfish sequence was used as the outgroup to root the tree.

### Orthogroup-based GO and KEGG enrichment analysis

Some orthogroups contain multiple members of the same gene family with highly similar functions; using all member genes directly for enrichment analysis would introduce family-biased results. We therefore constructed orthogroup-level GO and KEGG gene sets based on the well-annotated mouse GO/KEGG database. GO annotations were obtained from the org.Mm.eg.db v3.16.0, and KEGG annotations were derived from the mouse (mmu) pathway dataset of the KEGG database^111^.

Specifically, the GO/KEGG annotation of an orthogroup was defined as the union of annotations of all its member mouse genes. Hypergeometric test was performed with all annotated orthogroups as the background to calculate enrichment *P* values.

### Identification of mammalian association-related genes (MAGs)

The intermediate region of the Pr-Al gradient generally corresponds to the association cortex of each species. In mammals, association cortices are widely distributed, including prefrontal, temporal, posterior parietal and parts of the cingulate cortex. To more accurately identify genes associated with the association cortex in marmoset and mouse, we performed the analysis using regional-level whole-brain transcriptomic data^21^.

First, the Pr-Al index was converted to the Intersection Index, mapping the endpoints (Pr-Al index = ±1) to 0 and the mean of Pr-Al index to 1 with a smooth transition. The “cor.test” function from R was then used to calculate the Pearson correlation coefficient and *P* value between the expression of each gene and the Intersection Index. Orthogroups with *P* < 0.05 were defined as association-related genes. Finally, the intersection of association-related genes from marmoset and mouse was defined as Mammalian Association Genes (MAGs).

To examine the expression pattern of MAGs in other tetrapods, we scored z-score log normalized ST data of each species for the MAG set using Seurat’s “AddModuleScore”, and calculated the average MAG score per brain region. To ensure at least 3 biological replicates, the left and right hemispheres of different sections were treated as independent replicates. One-way ANOVA was performed using the “aov” function from R, followed by post-hoc *t*-tests for avian data using the t.test function.

To further validate the regional enrichment of MAGs, we performed K-means clustering of MAG expression across brain regions of 3 non-mammalian tetrapods (avian, turtle, axolotl) with *k* = 3. Genes highly expressed in avian M and MI regions were defined as mammal–avian association-related genes.

For subsequent Synaptic Gene Ontology (SynGO) analysis^77^, which does not accept orthogroup input, we used the full list of original genes corresponding to all mammal–avian association-related genes as input.

## Acknowledgements

We would like to thank DCS Cloud (https://cloud.stomics.tech/) for providing the computational resources and software support necessary for this study.

This Project was supported by China National Postdoctoral Program for Innovative Talents (BX20250145), National Natural Science Foundation of China (No. 32500590), and National Key Research and Development Program of China(2024YFF1206600).

Declaration of generative Al and Al-assisted technologies in the manuscript preparation process

We utilized a large language model for writing assistance and language polishing during manuscript drafting. All AI-modified text was carefully examined and finalized by the authors.

## Author contributions

S. H., Z. H, S. Li designed the methodology. Z. Z., D. C., X. D., Q. L., S. H. acquired the analysis resources. Z. Z., D. C., X. D., Q. L. performed the single-cell RNA-seq and Stereo-seq processing. Z. H, S. Li analyzed the study data. S. H., Z. H, S. Li, C. L., S. M., Y. S. wrote the manuscript. S. H. acquired the funding. S. H., Z. H, S. Li, C. L., S. M., Y. S. conceptualized the study. S. H., L. L., G. F., S. Liu supervised the project.

---

**Figure S1.**
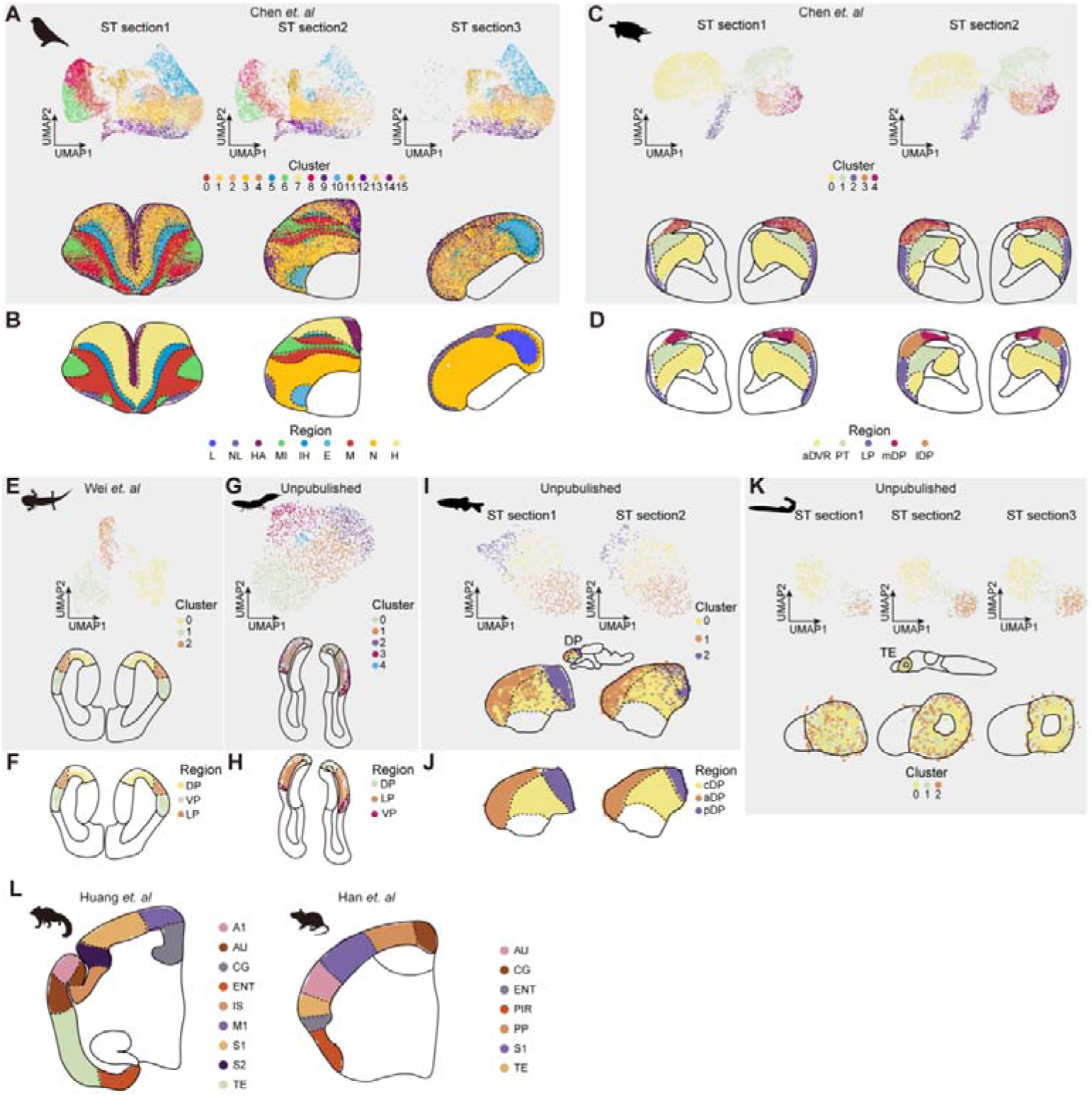
Pallial region definitions across all eight vertebrate species. (A, C, E, G, **I)** Unsupervised clustering of spatial transcriptomic (ST) sections from avian (A, 3 sections), turtle (C, 2 sections), axolotl (E, 2 sections), lungfish (G, 2 sections), and zebrafish (I, 2 sections); **(K)** Unsupervised clustering of 3 ST sections from lamprey, showing no spatially distinct cell clusters; **(B, D, F, H, J)** Manual region annotations of the corresponding ST sections shown in (A, C, E, G, I), respectively; **(L)** Schematic diagrams of standard brain atlases for marmoset (left) and mouse (right), illustrating the pallial regions included in this study. avian: L, field L; NL, nidopallium laterale; HA, hyperpallium apicale; IH, intercalated hyperpallium; E, entopallium; M, mesopallium; N, nidopallium; H, hyperpallium. Turtle: aDVR, anterior dorsal ventricular ridge; PT, pallial thickening; LP, lateral pallium; mDP, medial dorsal pallium; lDP, lateral dorsal pallium. Axolotl: DP, dorsal pallium; VP, ventral pallium; LP, lateral pallium. Lungfish: aDP, anterior dorsal pallium; cDP, central dorsal pallium; pDP, posterior dorsal pallium; VP, ventral pallium; LP, lateral pallium. Marmoset and mouse: A1, primary auditory cortex; AU, auditory cortex; CG, cingulate cortex; ENT, entorhinal; IS, insular; M1, primary motor cortex; PIR, piriform; PP, posterior parietal cortex; S1, primary somatosensory cortex; S2, secondary somatosensory cortex; TE, temporal.

**Figure S2.**
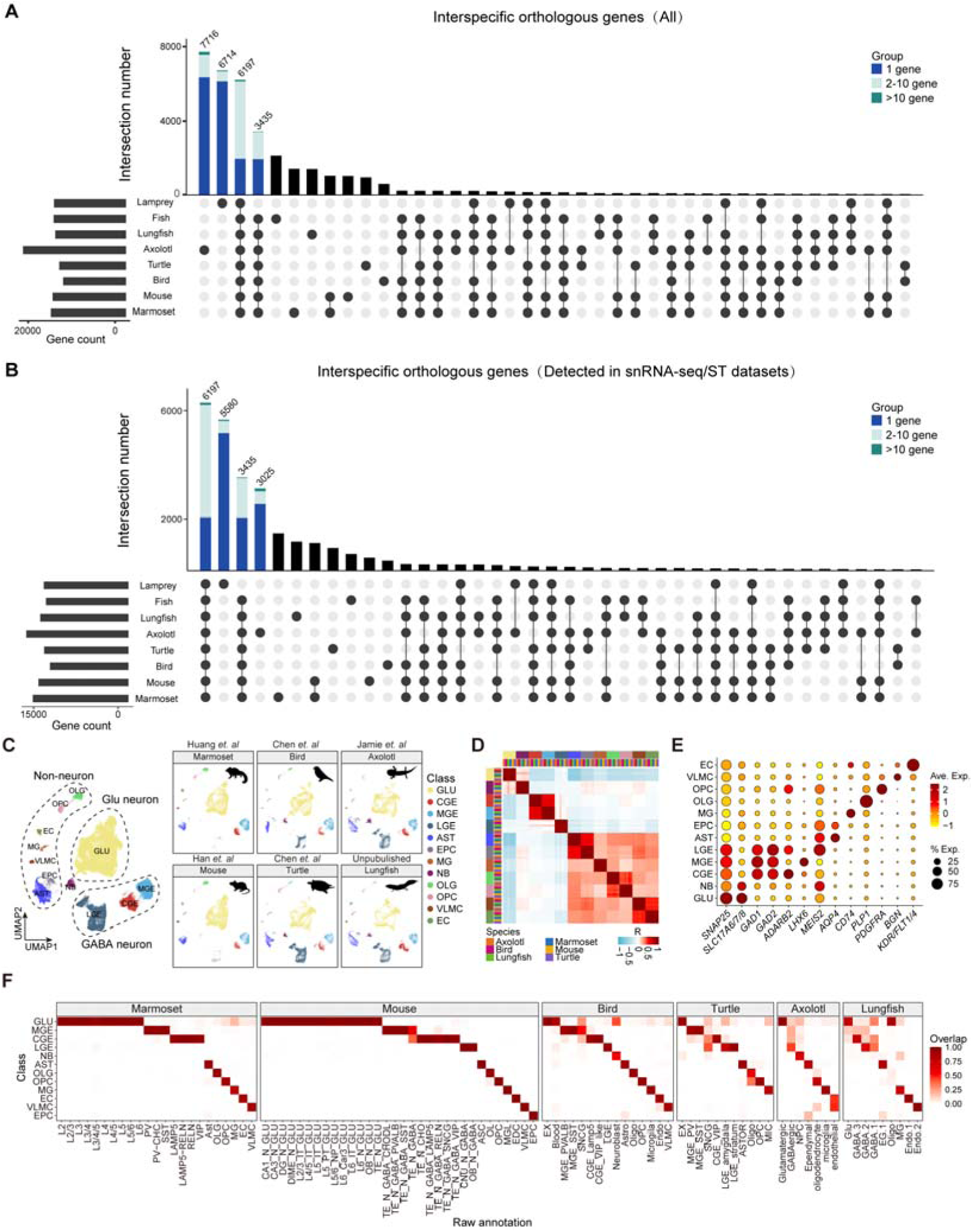
Cross-species orthogroup identification and integrated single-nucleus transcriptomic analysis. **(A)** UpSet plot showing the intersection of interspecific orthologous genes identified by OrthoFinder across all eight vertebrate species. Left bar plot indicates the total number of genes per species; top bar plot indicates the size of orthogroup intersections; dot-line matrix indicates the species composition of each intersection. **(B)** UpSet plot showing the intersection of orthologous genes detected in transcriptomic datasets. **(C)** UMAP visualization of integrated single-nucleus RNA-seq (snRNA-seq) data from six sarcopterygian species. Left panel shows three major cell classes (non-neuron, glutamatergic (Glu) neuron, GABAergic (GABA) neuron); right panels show UMAP projections for each individual species. **(D)** Pearson correlation (R) heatmap of homologous cell types across species. **(E)** Dot plot showing the expression of conserved cell type marker genes. Dot size indicates the percentage of cells expressing the gene; color intensity indicates average expression level. **(F)** Cell type overlap heatmap across species, showing the correspondence between original cell annotations and integrated unified cell types.

**Figure S3.**
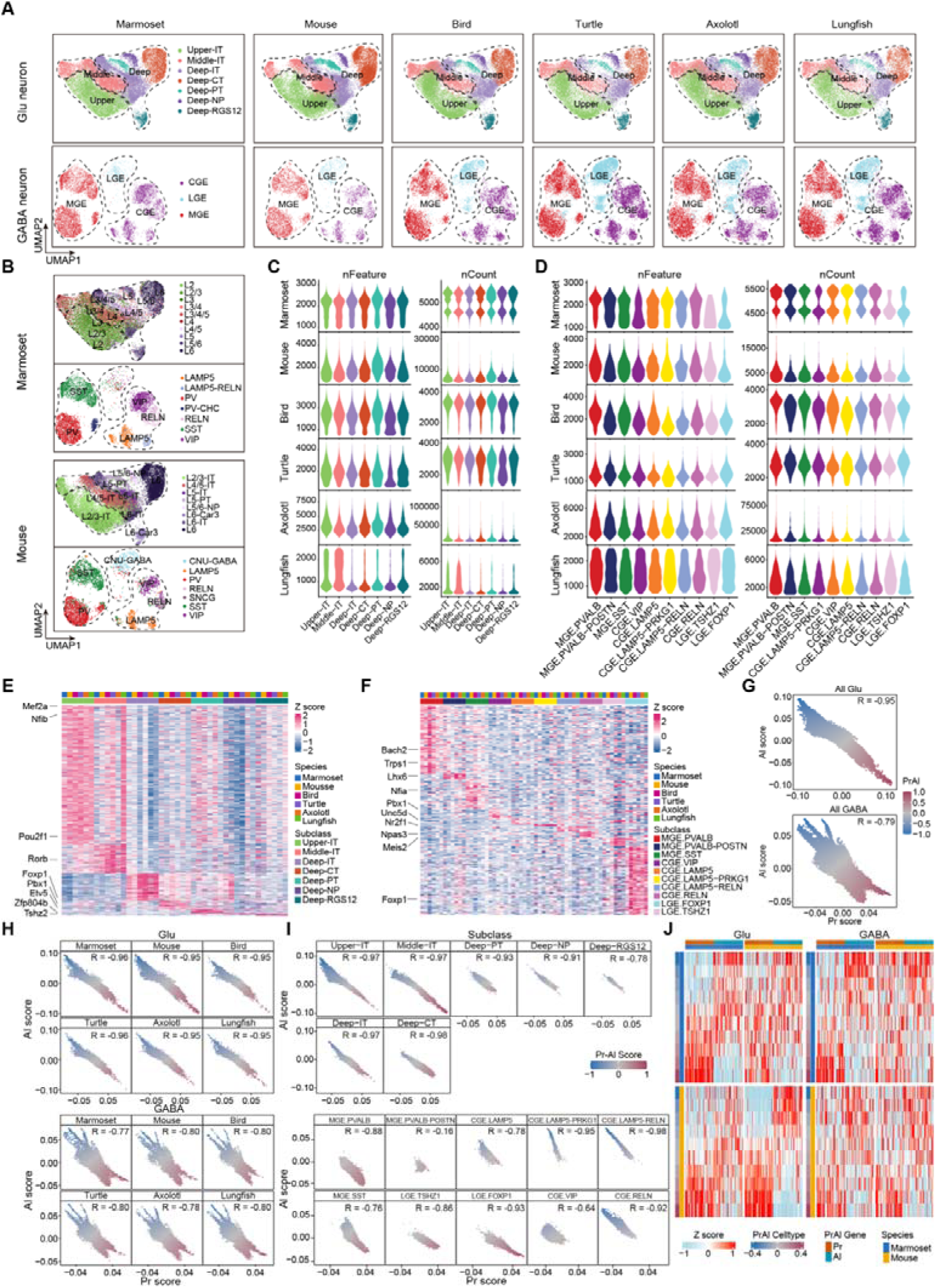
Validation of cross-species cell type integration and Pr-Al score analysis. **(A)** UMAP embeddings of integrated glutamatergic (Glu, top) and GABAergic (GABA, bottom) neurons across six species, colored by cell subclass. **(B)** UMAP visualizations of marmoset (top) and mouse (bottom) neurons show the distribution of distinct cell types, demonstrating the accuracy of cell subclass classification. **(C-D)** Violin plots showing the number of nFeature (left) and nCount (right) for Glu (C) and GABA (D) neurons across six species, stratified by cell subclass, confirming high-quality scRNA-seq data. **(E-F)** Heatmaps of top subclass marker genes defining Glu (E) and GABA (F) neurons, with expression scaled by Z-score, illustrating evolutionarily conserved cross-species expression patterns. **(G)** Scatter plots showing significant negative correlations between Pr- and Al-related gene set scores in integrated Glu (top; *R* = −0.95, *p* < 0.01) and GABA (bottom; *R* = −0.79, *p* < 0.01) neurons. **(H)** Multi-species scatter plots of Pr- and Al-related gene set scores in Glu (top) and GABA (bottom) neurons, stratified by species. **(I)** Correlation between Pr and Al gene set scores within each cell subclass, demonstrating consistent negative correlations across all subclasses. **(J)** Heatmap of z-score normalized expression of individual top 200 Pr- and Al-related genes, confirming opposing expression patterns between Pr- and Al-related genes across cell types binned by Pr-Al scores in marmoset and mouse neurons.

**Figure S4.**
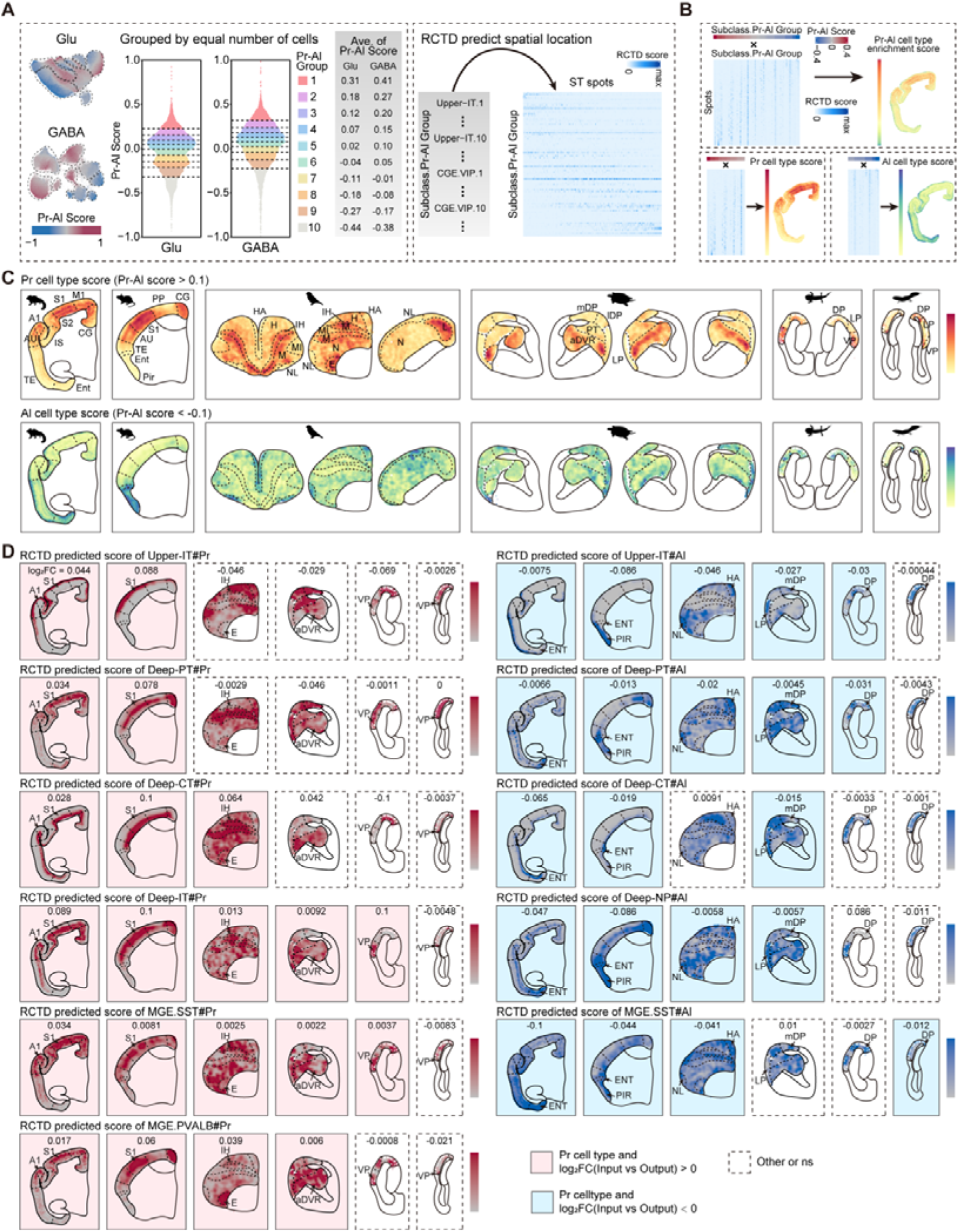
Spatial deconvolution analysis of Pr-Al gradient cell types across sarcopterygians. **(A)** Left, UMAP visualization of integrated glutamatergic (Glu) and GABAergic (GABA) neurons colored by Pr-Al score. Violin plots show the distribution of Pr-Al scores in Glu and GABA neurons, which were divided into 10 equal-sized groups. Right, table showing the average Pr-Al score for each group, with representative gradient-resolved cell types used for RCTD spatial deconvolution labeled. **(B)** Workflow and example results of RCTD spatial deconvolution analysis. Top, heatmaps showing subclass Pr-Al group scores and corresponding RCTD predicted scores in ST spots. Bottom, spatial maps showing the distribution of Pr or Al cell type scores and all cell type scores. **(C)** Spatial distribution of aggregated Pr cell type scores (Pr-Al score > 0.1, top) and Al cell type scores (Pr-Al score < −0.1, bottom) across all six sarcopterygian species. **(D)** Spatial distribution of RCTD predicted scores for representative Pr-enriched (left) and Al-enriched (right) cell types across species. Numbers indicate log2 fold change (log2FC) of cell type enrichment between input and output regions. Red shading indicates Pr subtypes significantly enriched in input regions (log2FC > 0); blue shading indicates Al subtypes significantly enriched in output regions (log2FC < 0); white indicates no significant difference (ns).

**Figure S5.**
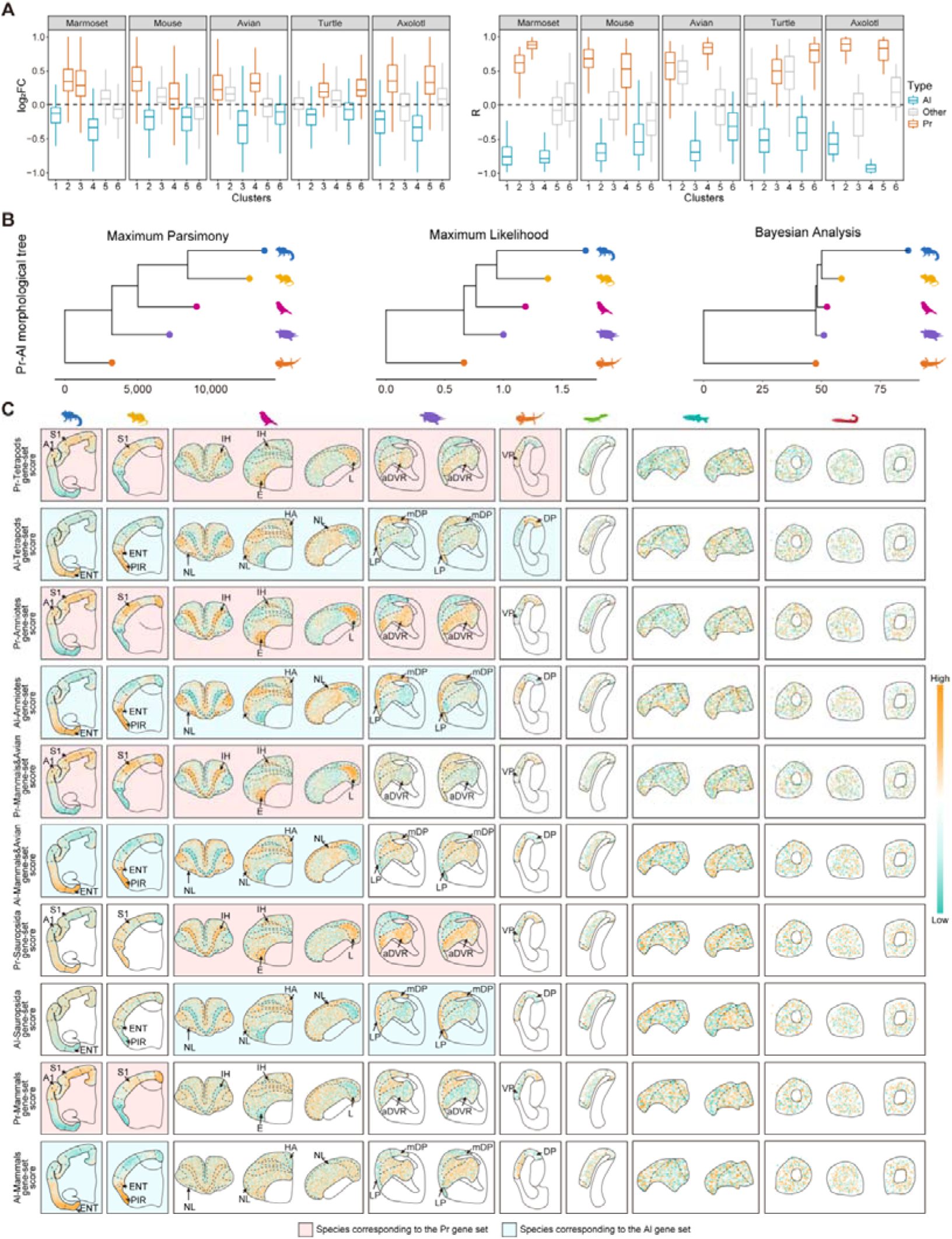
Quantitative validation of Pr-Al gene cluster definition and evolutionary conservation in single-cell data. **(A)** Quantitative metrics for defining Pr-enriched and Al-enriched gene clusters from Fig. 4A. Log2 fold change (log2FC, left) of gene expression between input and output regions across five tetrapod species. Pearson correlation coefficients (R, right) between gene expression levels and the Pr-Al regional order (from Al-dominant to Pr-dominant) across five tetrapod species. **(B)** Tree constructed from Pr-Al gene sets across species using the maximum parsimony (left), maximum likelihood (middle) and bayesian analysis (right) methods. **(C)** Spatial distribution of evolutionarily stratified Pr-Al gene set scores across all eight vertebrate species. Rows correspond to gene sets, and columns correspond to species. Red and blue shading indicates the species corresponding to the Pr and Al gene set, respectively.

**Figure S6.**
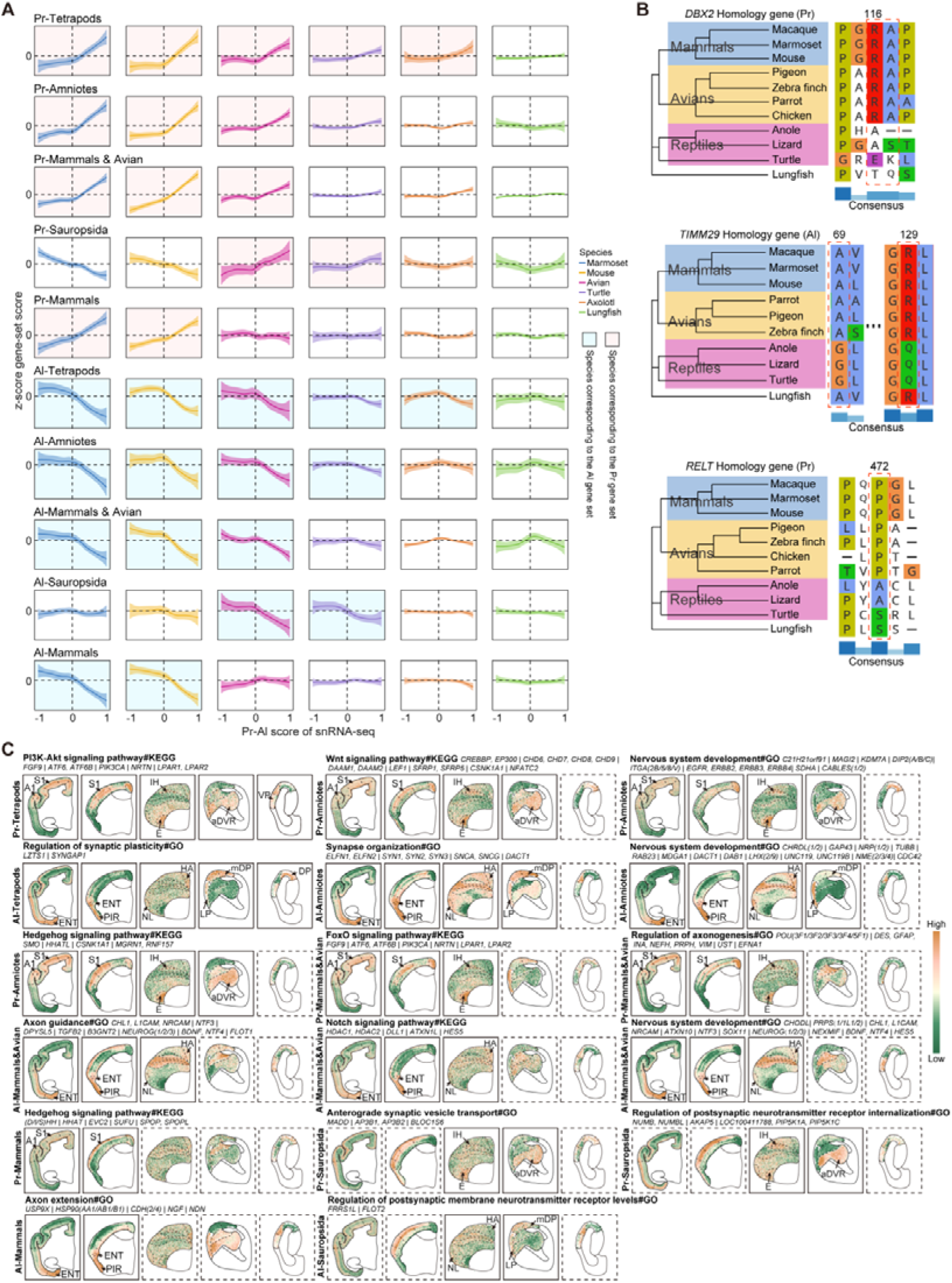
Spatial validation of evolutionarily stratified Pr-Al gene sets and functional pathways. **(A)** Single-cell transcriptomic validation of the evolutionarily stratified Pr-Al gene sets from Fig. 4C. Red and blue shading indicates the species corresponding to the Pr and Al gene set, respectively. **(B)** Protein sequence alignments of three representative mammals-avian conserved Pr or Al genes, showing lineage-specific amino acid substitutions: *DBX2* (Pr), *TIMM29* (Al), and *RELT* (Pr). **(C)** Spatial distribution of gene set scores for representative enriched Gene Ontology (GO) and KEGG pathways from Fig. 4G. Arrows indicate Pr or Al regions with significant pathway enrichment.

**Figure S7.**
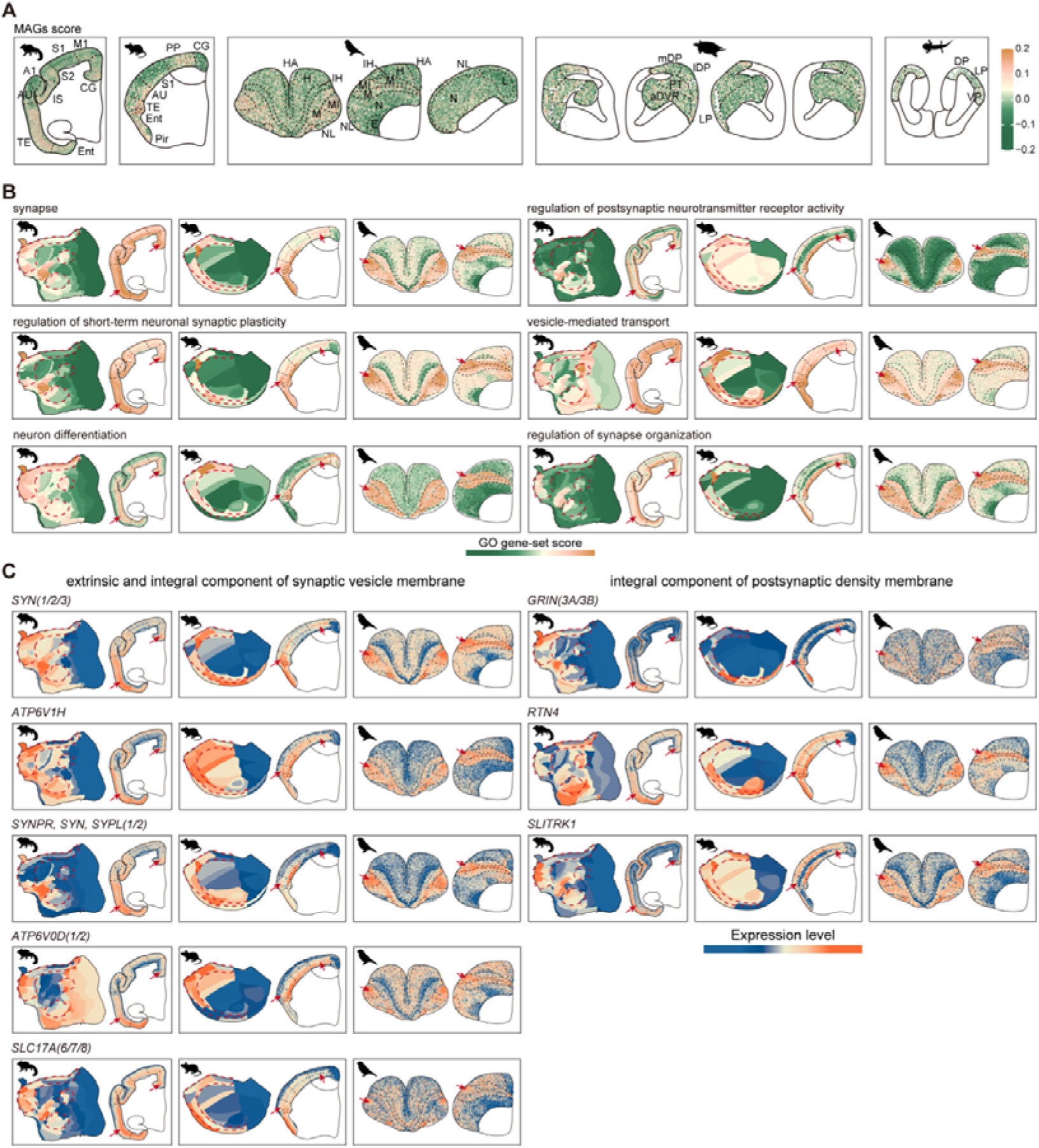
Spatial validation of mammalian association genes (MAGs) and their functional gene sets across species. **(A)** Spatial distribution of MAGs scores across all six sarcopterygian species (from left to right: marmoset, mouse, avian, turtle, axolotl, lungfish). **(B)** Spatial distribution of gene set scores for the six significantly enriched Gene Ontology (GO) terms related neuronal function from the convergent mammals-avian association gene module. Red dashed lines and arrows delineate association cortex regions. **(C)** Spatial expression patterns of representative synaptic genes from the SynGO.

